# Empirical and Computational Comparison of Alternative Therapeutic Exon Skip Repairs for Duchenne Muscular Dystrophy

**DOI:** 10.1101/527705

**Authors:** Krystal Manyuan Ma, Evelyn S Thomas, Jeff Wereszczynski, Nick Menhart

## Abstract

Duchenne muscular dystrophy is a common and devastating genetic disease that is primarily caused by exon deletions that create a genetic frameshift in dystrophin. Exon skipping therapy seeks to correct this by masking an exon during the mRNA maturation process, which restores dystrophin expression, but creates an edited protein missing both the original defect and the therapeutically skipped region. Crucially, it is possible to correct many defects in alternative ways, by skipping an exon either before, or after the patient’s defect. This results in alternatively edited, hybrid proteins, of possibly different properties and therapeutic consequences. Here, we examined three such dystrophin exon skipped edits, comprising two pairs of alternative repairs of the same underlying DMD defect. We found that in both cases, one member of each alternative repair was more stable than the other by a variety of thermodynamic and biochemical measures. We also examined the origin of these differences by molecular dynamics simulations, which showed that these stability differences were the result of different types of structural perturbations. For example, in one edit there was partial unfolding at the edit site which caused domain-localized perturbations, while in another there was unfolding at the protein domain junctions distal to the edit site which increased molecular flexibility. These results demonstrate that alternative exon skip repairs of the same underlying defect can have very different consequences at the level of protein structure and stability, and furthermore that these can arise by different mechanisms, either locally, or by more subtle long-range perturbations.

## Introduction

Duchenne muscular dystrophy (DMD) is a severe and unfortunately prevalent (~1:5000 male births) pediatric genetic disease caused by the loss of dystrophin proteins from muscle cells (Birnkrant et al., 2018; Falzarano et al., 2015; Hamuro et al., 2017). The prognosis for those with this condition is bleak: progressive muscle wasting leading to loss of ambulation in the second decade of life, with death typically in the second or third decade of life. Exon skipping is the first, and so far, only curative therapy for this condition, and offers the promise of a significantly better life to these patients and their families.

The most common type of genetic defect underlying DMD is a large (~1 to 100kbp) deletion of one or more exons. In a protein as large as dystrophin (~427 kDa) this often results in the juxtaposition of exons of incompatible reading frames and also inevitably introduces a new stop codon which truncates the protein. The nonsense mediated decay system generally flags and destroys these defective transcripts (Khajavi et al., 2006; Linde and Kerem, 2008) so that even this truncated protein is not expressed at anything approaching normal levels, and dystrophin is typically undetectable in DMD cells.

Exon skipping seeks to address this deficiency by masking additional exons to correct the reading frame and restore dystrophin expression. Clinically, this is done with antisense oligonucleotide analogs, AONs, which are similar to short, ~20 bp DNA primers, but with modified backbones to improve pharmacological properties (Aartsma-Rus et al., 2017; Nakamura, 2017). This aims to restore expression by correcting the damaged reading frame, with the clinical aim of slowing or reversing muscle deterioration. However, although reading frame correction restores protein expression, the proteins produced lack the regions corresponding to the patient’s original defect, as well as the therapeutically skipped exon, thus altering the protein’s structure. While deleting a region corresponding to several exons from a protein’s structure and reattaching the remaining regions (which are not normally proximal, and so have not evolved to be structurally compatible) may seem like a drastic alteration, we have shown that in some cases stable structures can form (Menhart, 2006).

Dystrophin acts largely as a mechanical stabilizing agent (Harper et al., 2002; Le et al., 2018) and is thought to work as a molecular shock absorber, so changes in its structure and stability likely alter those functions. In addition, dystrophin also contains a variety of binding sites for other proteins – most significantly neuronal nitric oxide synthetase (Lai et al., 2009) and phospholipids (Sarkis et al., 2011) – and edits may disrupt and modify those interactions (Sahni et al., 2012). How edits impact the structure and function of dystrophin is an open question and presumably of clinical importance. Insight into this is provided by Becker’s Muscular dystrophy (BMD) which is caused by naturally occurring in-frame defects in dystrophin. BMD is a milder but more heterogeneous condition, varying from nearly as severe as DMD, to nearly benign – with the suspicion that some defects might be so benign as to be sub-clinical (Le Rumeur, 2015; Zimowski et al., 2017). Exon skipping essentially aims to convert DMD to therapeutically induced BMD. However, due to the complexity of the dystrophin gene, there are a very large number of defects known. While in aggregate, many edits have been identified, many individual defects are known only from a few cases – or are unknown. It thus becomes difficult to accurately assess clinical impact, as patients with the same defect can progress very differently. However, by pooling many different types of edits, many studies have shown a clear and significant relationship between edit type and clinical severity (Anthony et al., 2014; Findlay et al., 2015a; Kaspar et al., 2009a) – we just are not able to accurately compare arbitrary pairs of defects in most cases.

Making these types of comparisons is becoming increasingly important as additional exon skipping compounds become available. In many cases a patient’s underlying defect can be treated in different ways, skipping different exons, to create alternative repairs with potentially different properties and clinical outcomes. Currently there is only one choice for exon skipping, eteplirsen (Exondys51) which targets exon 51 (Aartsma-Rus and Krieg, 2017). However clinical trials for compounds targeting exons 44, 45, and 53 are underway (e.g. NCT02329769, NCT02500381, NCT01957059 in the US, and other programs worldwide (Komaki et al., 2018; Nakamura, 2017; Shimizu-Motohashi et al., 2016)), and preclinical programs exist for many other exons. If development of the therapies targeting these new exons are successful, the prospect of patients and their physicians facing a choice about which to use will become a reality. Since different therapies will result in the expression of distinct dystrophin structures, a rational, structure-based basis for making this choice is needed.

DMD causing mutations are most commonly found in two hotspot regions of the gene since this is where exons of different starting and ending reading frames are clustered. These occur in the central rod region of the protein, which consists of 24 of copies of a rod domain known as spectrin type repeat (STR) (Djinovic-Carugo et al., 2002; Nicolas et al., 2014). The largest and most significant of these is the hotspot 2 region, spanning exons 43 to exon 59, which corresponds to STRs D16 to D24 (Figure 1). Within this region there are 64 possible exon-edited proteins. While the structures of all of these edited proteins are interesting and applicable to DMD therapy, here we are most concerned with pairs of edits that may be end points of alternative exon skip repairs of the same underlying DMD mutation. These cases represent those where a choice of therapy exists, and are more urgently needed in the near term as exon skipping choices become available. The most common type of choice occurs when either the exon immediately before, or immediately after, the original deletion is targeted. For instance, patients with an out-of-frame exon 52 defect could be treated to either skip exon 51 with eteplirsen to create an in-frame Δe51-52 skip; or a potential exon 53 skip reagent creating Δe52-53. Not every defect has such simple alternatives, but many do: such pairs lie adjacent on a diagonal from top left to bottom right in Figure 1, and there are 31 such edits. More possibilities arise with multi-exon skipping which is a more demanding task but has been studied in cell culture and animal models. Furthermore, in general terms, the question of *“what edit to make”* is of even greater importance with more flexible techniques, such as CRISPR gene editing, which although not yet in the clinic, is widely viewed with great promise and potential.

**Figure 1.**
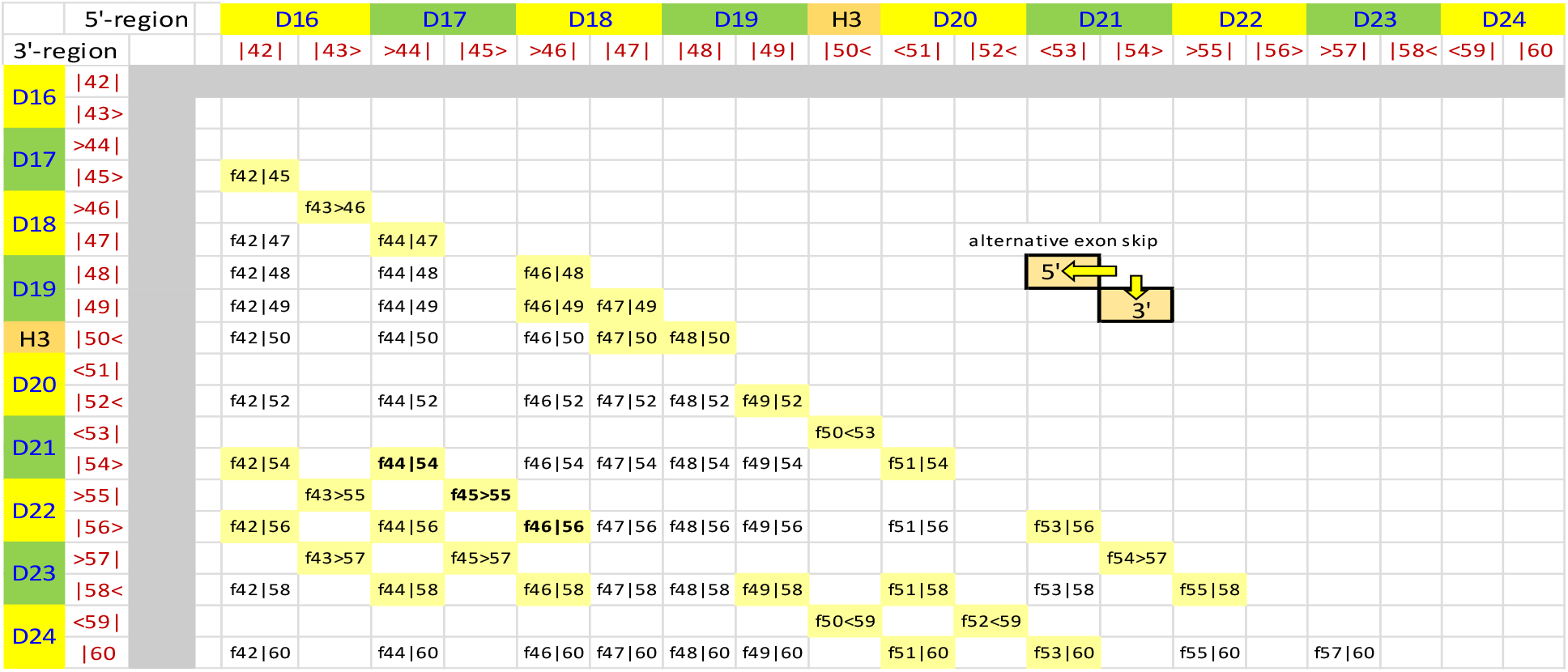
All possible exon edits in the Hotspot 2 region, exon 42 to 60. Viable exon edits result in in-frame fusion if the ending phase of the 5’ remaining exon is compatible with the starting phase of the 3’ remaining exon. There are 3 possible reading frames for each exon boundary { <, |, >}corresponding to the protein reading frame at the exon junctions {-1 or one base before the end of a codon, 0 or at the codon boundary, +1 or one base into a codon}. The approximate STR boundaries are also shown in alternating green and yellow, although it must be emphasized that STR boundaries do not exactly correspond with exon boundaries. There are 64 in-frame fusions (names as: f45>55 is a deletion of 46 to 54 fusing exons 45 and 55 at a phase 1 boundary). Blank cells are unviable, out-of-frame DMD defects. Alternative repairs occur when skipping of either the 5’ or 3’ exon converts a DMD defect into an in-frame edit. These lie in pairs along a diagonal as shown. Of the 64 possible edits, 31 share this property. We selected 3 edits that comprise two alternative repairs: f44|54, f44>55, and f45|65, shown in bold and more conventionally referred to by the exons deleted, i.e. Δe45-53, Δe46-54, and Δe47-55.

Since there are many possible edits, with more when the entire dystrophin gene is considered, and even more when other editing modalities (CRISPR etc.) might be possible in the future, we are conducting studies to determine not only *“which edit is best*?”, but also to attempt to determine the basic principles that make an edit better or worse. This obviously involves studies at a range of scales from atomic to organismal, but we are focusing in the impact on the protein structure, since it is the dystrophin protein that performs the crucial muscle stabilizing function. Here, we selected one such set of three edits, Δe45-53, Δe46-54, and Δe47-55, encompassing alternative exon skipping repairs of two frameshifting DMD-type deletions: Δe46-53 and Δe47-54 (Table 1) and demonstrated that one of these, Δe46-54, produced a protein that is significantly more stable than the other two Δe45-53 and Δe47-55. Furthermore, we conducted molecular modelling and molecular dynamics (MD) simulations on these edits to elucidate the molecular basis of these differences. We found that the destabilizing edits did not appear to cause structural perturbations proximal to the edit site, but rather function distally, at STR junctions that became “non-wild-type” when regions away from the edit site that are not normally in contact are spatially juxtaposed. This provides the first molecular level understanding of these stability differences in exon edited rods.

**Table 1.** Exon Repairs in this Study. Edits in this study were chosen to be alternative repairs of specific DMD type defects

| DMD type defect (out of frame) | exon skip target | in-frame repair edit studied |
| --- | --- | --- |
| $\Delta e46-53$ | exon 45 | $\Delta e45-53$ |
| | exon 54 | $\Delta e46-54$ |
| $\Delta e47-54$ | exon 46 | |
| | exon 55 | $\Delta e47-55$ |

## Methods

Concise methods are given, with complete details available in Supplementary Methods. The dystrophin protein studied has UniProt ID P11532.

### Target Selection

In order to most easily study the local perturbations introduced by exon skipping, we produced exon edited targets in the context of as small a region as possible that has native-like dystrophin rod ends (Findlay et al., 2015b; Nicolas et al., 2015). This entails identification of biophysical domain boundaries of the dystrophin rod to locate appropriate termini. In prior work(Mirza et al., 2010), we determined that D16 and D17 form a thermodynamically cooperating domain, as do D20 to D22. Referring to Figure 2, boundaries at N-terminal to D16 and C-terminal to D22 flank all the edits sites and might be suitable. However, one of our edits, Δe47-55 removes large portions of D22 (53 of the 117 amino acids, 45%) leaving behind a fractional STR at the C-terminus. To ensure well-formed rod ends, we decided to in addition utilize a second cohort of experimental targets with a C-terminus boundary at D24. As such we produced two series of exon edited proteins, in both D16:22 and D16:24 parent molecules.

**Figure 2.**
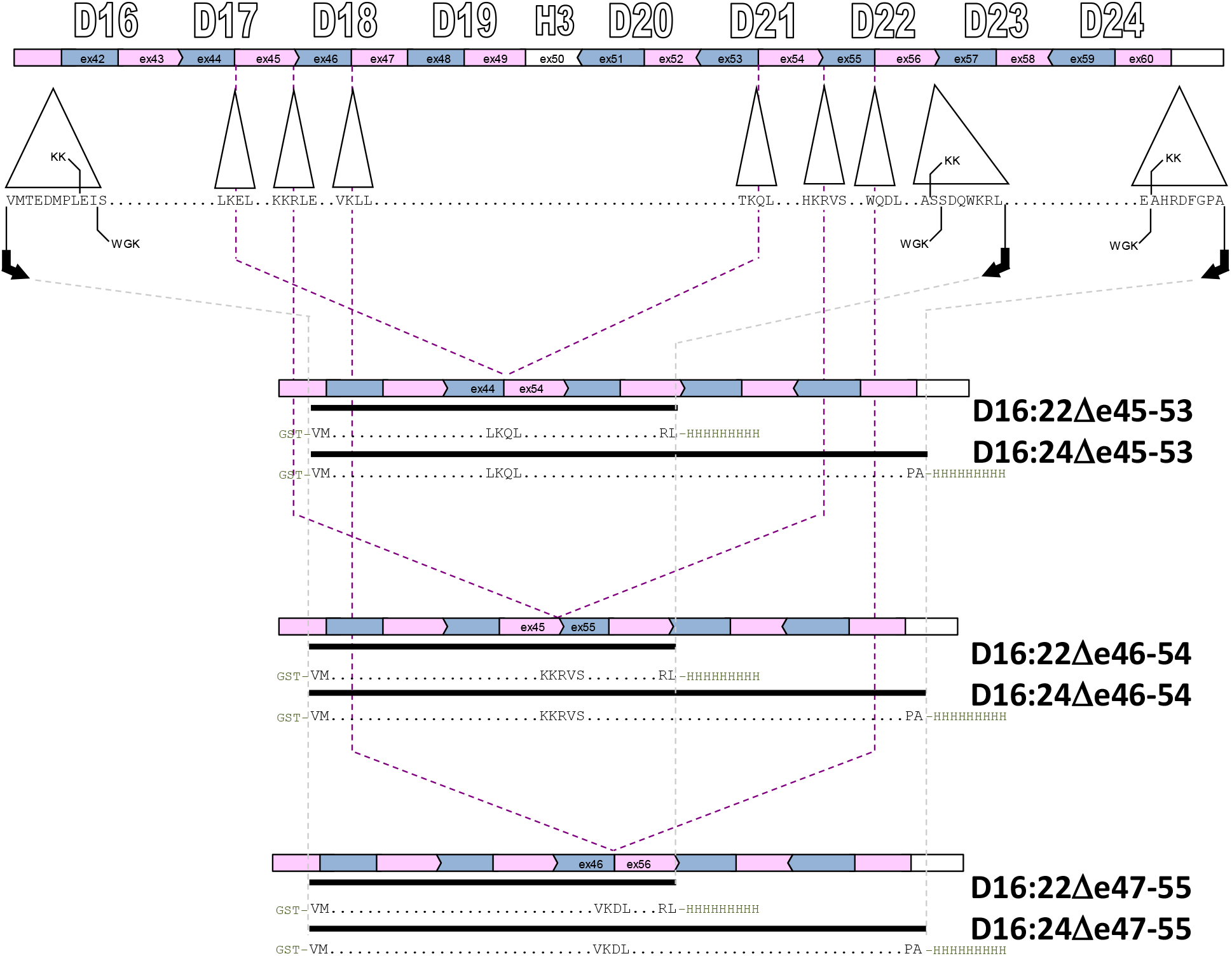
Cloning strategy. Target proteins were expressed in both D16:22 and D16:24 parent molecules. At the top, the exon structure is shown in alternating pink and blue bars, aligned with the STR structure in this region. There are two alternative alignments describing the endpoints of dystrophin STRs as described in the text, KK and WGK. These differ by at most 8 amino acids, so we utilized endpoints that are 8 amino acids larger than the KK alignment to be compatible with both, as in our previous work on wild type and exon edited STRs. PCR and Gibson assembly were used to achieve the exon edits (indicated by the dashed lines) to add glutathione-S-transferase, GST and His9 affinity tags.

### Cloning and Protein Production

These proteins were cloned, expressed, and purified as previously described (Sahni et al., 2012). Briefly, plasmids containing the genes coding for the target proteins were assembled by the Gibson protocol (Gibson et al., 2010) using unedited dystrophin genes fragments previously studied, and transformed into *E. coli* NEB express (a commercially modified DH5α strain optimized for protein expression). Concordance of all constructed targets with the intended sequences as indicated in Figure 2 was confirmed by direct DNA sequencing over the entire dystrophin region.

These proteins were expressed in a double affinity-tagged form utilizing an N-terminal GST (glutathione S-transferase) domain and a C-terminal His9 tag. Only the full-length, undegraded protein was selected by both affinity protocols. To remove any remaining non-full-length species, a final ion exchange purification step was used. All proteins were purified to a single band > 98% at the appropriate molecule weight as assessed by SDS-PAGE.

#### Biophysical Characterization

These proteins were characterized as previously described (Sahni et al., 2012). Since STRs are triple α-helical motifs, we used total helicity by circular dichroism, CD, to assess how well-folded they were initially, reported as both mean residue ellipticity at 222 nm, ϕ_222_; as well as fractional helicity as determined by a simple but robust 3-basis set model (Greenfield and Fasman, 1969) that has been extensively used by us and others (Calvert et al., 1996; Ipsaro et al., 2008; Wolny et al., 2011) for helicity of STR proteins. To measure overall thermodynamic stability, we performed thermal denaturation as followed by CD, which provided the melting temperature, Tm, as well as the enthalpy of unfolding, ΔH. Here, we used the signal at 222 nm, ϕ_222_, which was collected every 0.2 C and fast Fourier transform (FFT) filter smoothed (lowpass cutoff 0.2 C) and then the derivative, dϕ_222_/dT fit to the John-Weeks equation to yield ΔH and Tm (John and Weeks, 2000). Because unstructured regions are invisible to denaturation sensitive assays, we used sensitivity to protease challenge to assess how much and to what extent the target protein contained poorly folded regions (Fontana et al., 1997; Hubbard, 1998). To avoid primary sequence bias, we use a highly non-specific protease, Proteinase K, PK. The specific measure we report is the concentration of PK that achieves 50% degradation in a defined assay (30 min at 37 °C in PBS buffer), the PK_50_ value. This specific assay has been applied by us and others (McCourt et al., 2015) to a wide range of native and exon edited dystrophin rods.

Our standard of reproducibility was that all data was acquired from at least three independent experiments on at least two independent batches of protein. Our standard of statistical significance is P<0.05 (significant) or P<0.005 (highly significant) using a two tailed Student’s t-test, as well as an effect size d>2, where d is Cohen’s 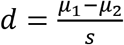.

#### Computational Modeling

##### Initial Model Development

Since empirical studies of the three target edits produced similar results in both the D16:22 and D16:24 contexts, only the D16:22 case was pushed forward to computational studies due to its smaller size. The primary sequences of three targets in the D16:22 family were submitted to the Robetta automated structure prediction server (Kim et al., 2004). The output models were ranked by combination of ProQ2 score (Ray et al., 2012) and TM-score (Zhang and Skolnick, 2004), but all were grossly similar in that all were rod-like, STR containing structures.

##### Implicit Solvent Molecular Dynamics Simulations

The top model for each target was chosen for further computational study. These structures were processed to removed hydrogens in VMD (Humphrey et al., 1996) and then the topology and coordinate files were developed using the LEaP package of the AMBER software suite (Case et al., 2005). Structures were minimized and equilibrated through a multi-stage protocol (more details can be found in Supplemental Methods). We conducted six independent MD production runs (with independent equilibration), three of 2 μs; and three shorter runs of 0.5 μs, with the first 250 μs discarded as further equilibration. That 250 μs was sufficient equilibration time was justified by examining both total helicity and backbone RMSD, Figure S1, for lack of system drift after that time. Conformations after that were saved every 50 ps, for a total of 120,000 coordinate sets for each target, which was the standard implicit dataset. All implicit solvent calculations utilized the “GB-Neck2” solvation model and parameters developed by Nguyen, Roe, and Simmerling, which was enabled with the igb = 8 AMBER configuration keyword and the mbondi3 radii set (Nguyen et al., 2013). The SHAKE algorithm was used with a 2 femtosecond timestep and the AMBER ff14SB forcefield (Maier et al., 2015), along with the GPU accelerated version of PMEMD (Götz et al., 2012). Simulations were performed on local resources, as well as the Extreme Science and Engineering Development Environment (XSEDE) (Towns et al., 2014).

##### Explicit Solvent Molecular Dynamics Simulations

These initial datasets were developed using an implicit solvent model, which is a tradeoff between physical concordance and computational tractability. To examine this and provide reassurance that solvation effects were not unduly perturbing our data, an additional set of simulations was performed in an explicit solvent environment. As starting points for explicit simulation, the implicit standard dataset for each target was clustered using the hierarchical agglomerative algorithm (Shao et al., 2007) with average-linkage. Representative structures from all clusters with >1.5% abundance (four for Δe45-53 and Δe46-54 and three for Δe47-55) were solvated at 150 mM NaCl and pH 7, relaxed by a multistep protocol and subject to three 250 ns explicit MD simulations. Only the last 150 ns were used for analysis, (i.e. 450 ns for the three runs) and conformations were once again saved every 50 ps, for a total of 9,000 coordinate sets for these three runs from each starting structure. This yielded a standard explicit dataset of 36,000 coordinate sets for the four clusters run of Δe45-53 and Δe46-54; and 27,000 for coordinate sets for the three clusters run of Δe47-55.

##### Computational Analysis

Standard datasets obtained above were analyzed for helicity using cpptraj (Roe and Cheatham, 2013) which itself uses the DSSP algorithm which detects an (i,i+4) hydrogen bond (Kabsch and Sander, 1983). Inter-STR bending was assessed by defining an STR vector from the positions of conserved heptad hydrophobic residues (see Figures 6 and S2, as well as Supplementary Methods) in each STR, and then the angle between vectors of adjacent STRs was calculated with cpptraj. Pairwise intra-residue interaction energies within each STR were calculated by the Molecular Mechanics/Generalized Born Surface Area (MMGBSA) protocol(Chowdhury et al., 2013), with Generalized Born implicit solvent model (Mongan et al., 2007) using the MMPBSA.py script(Miller et al., 2012). Coordinated motions indicative of structural interactions were assessed by general motional correlation(Lange and Grubmüller, 2006). Visualization was performed in VMD and modified in GIMP (Peck, 2008), and data plots were created by OriginLab and Matplotlib (Hunter, 2007).

## Results

### Experimental

#### Secondary Structure by CD

All proteins exhibited circular dichroism (CD) spectra with double minima at ~208 nm and ~222 nm, which is characteristic of highly α-helical proteins (Figure 3). However, they varied significantly in either ϕ_222_ or the fractional α-helix content, fα, with Δe46-54 having the highest values, Δe47-55 close behind and Δe45-53 the lowest. These were consistent in both D16:22 and D16:24 parents. Crystal structures of non-dystrophin STRs show helicity values in the 85-90 % range, and spectroscopic studies of unskipped dystrophin 2-STR rods in this region show values in the 65-87 % range (D16:17 69 %, D17:18 69 %, D20:21 77 %, D21:22 87 %, D23:24 65 %, (Mirza et al., 2010)). By this measure, both Δe46-54 and Δe47-55 exhibit unremarkable helicity values consistent with typical STRs. On the other hand, Δe45-53 exhibits significantly lower helicity values, 36 % in D16:22 and 49 % in D16:24 that are not consistent with a well formed STR type structure for all of this protein, and suggest a significant disruption in structure for this edit.

**Figure 3.**
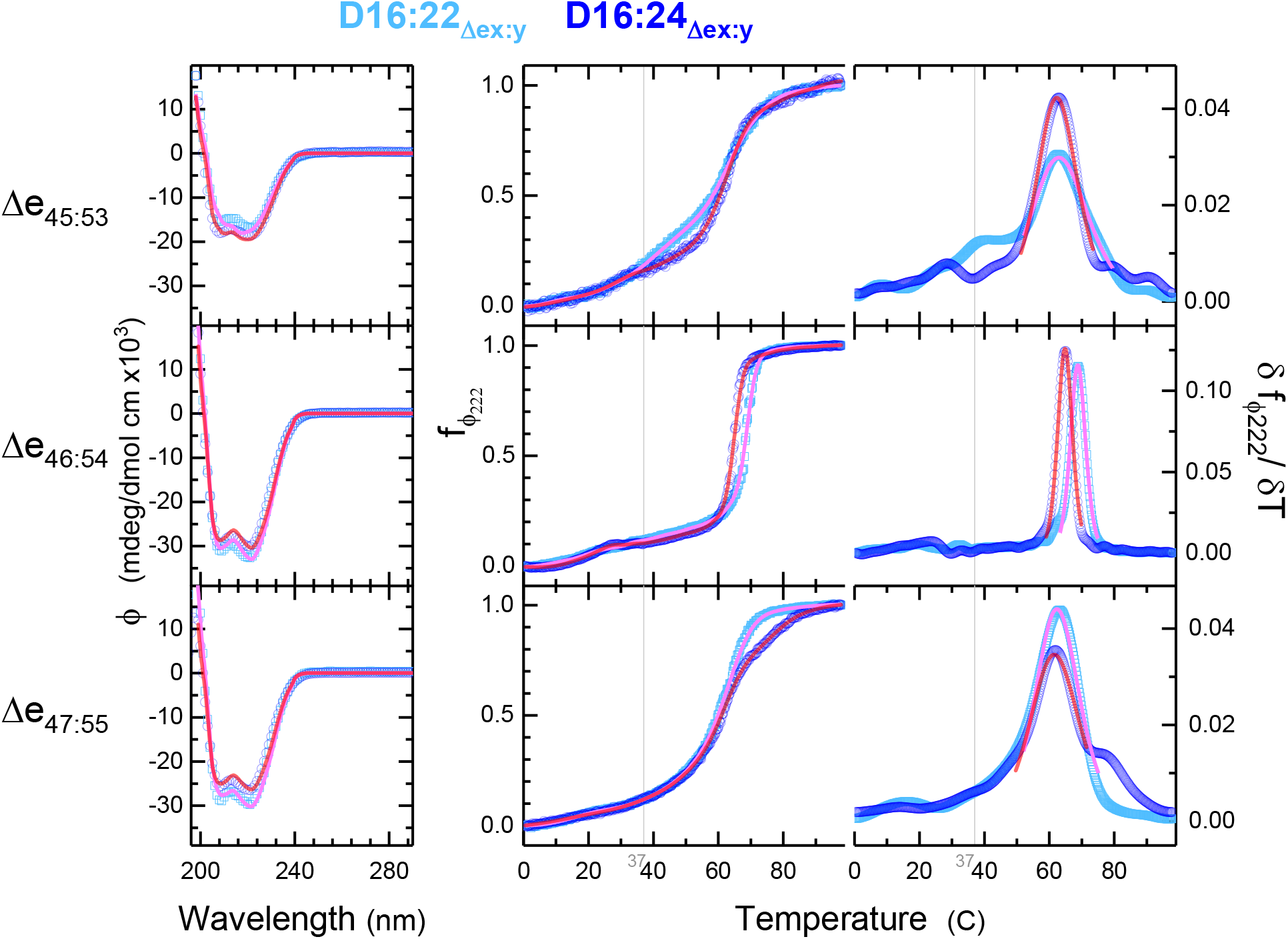
CD Structure and Stability. Analysis CD and thermal denaturation yield secondary structure and unfolding energies as measures of stability. Left is CD spectra at 25C, with cyan showing D16:22 and blue D16:24 parents. To the right is shown raw thermal unfolding profiles and the derivative plots for each system for temperature range of 0.5-98.5 C. Fit curves in pink, D16:22, and red, D16:24, show the three-component secondary structure analysis of the CD spectra. Fit curves in the raw thermal denaturation profiles, center, indicate the FFT filtered data; and on the derivative plot, right, the thermodynamic fit used to determine Tm and ΔH. The grey vertical line is at the physiologically relevant temperature of 37 C.

#### Thermodynamic Stability by Thermal Denaturation

Structural stability was also probed by thermal denaturation as monitored by circular dichroism. The three edits all displayed typical sigmoidal unfolding transitions, in the 50-70 °C region (Figure 3). By taking the derivative of the signal with respect to temperature, we can obtain thermodynamic properties of these transitions more accurately, and crucially, largely independent of spectroscopic parameters (John and Weeks, 2000). This yields precise melting temperatures in the 62-64 °C range for all edits. Once again, Δe46-54 was the most stable, but only by a few degrees. All these are in the range of typical STRs in this region (D16:17 69 °C, D17:18 55 °C, D20:21 69 °C, D21:22 70 °C, D23:24 64 °C (Mirza et al., 2010)).

However, when we examine how sharp the transitions are, we see dramatic differences in their widths, most evident in the derivatives of the plots on the right side of Figure 3. This width is most strongly impacted by the enthalpy of unfolding, ΔH, via the van’t Hoff equation 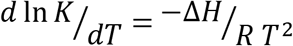 (which is implicit in the slightly more complex fitting equation described in the Methods), with narrow transitions resulting from large enthalpies and broad transition small enthalpies. We observe that Δe46-54 has a significantly narrower transition than either Δe45-53 or Δe47-55 indicating significantly different ΔH. A full analysis as described provides an unfolding enthalpy of 565 or 624 kJ/mol (in the D16:22 and D16:24 contexts respectively, see Table 2), which is consistent with – and even a bit larger than – unedited STR values in this region. Here, comparison to wild type, unedited dystrophin is a bit complex since the full unedited parent is a repetitive, multidomain protein that will have many complex overlapping transitions. However, we have previously demonstrated that these complex transitions can be effectively modelled as the sum of 2-STR transitions (Mirza et al., 2010), and also characterized all 2-STR transitions in the intact molecule. The wild type STR regions involved in this study vary from 304 to 539 kJ/mol (D16:17 504 kJ/mol; D17:18 340 kJ/mol; D20:21 529 kJ/mol; D21:22 531 kJ/mol, D23:24 345 kJ/mol (Mirza et al., 2010)). The other two edits, Δe47-55 and Δe45-53 exhibit broad low enthalpy transitions, in the 163-215 kJ/mol range, which is less than these values for typical STRs in that region.

Importantly, with exon skipping therapy patients do not have the option to move back to wild type, but rather to choose between two alternative edits. As such, the most relevant comparison is to compare alternatives to each other, not to wild-type. In our data analysis, we judged the statistical relevance of these comparisons by a two-tailed Student’s t-test, and in this case Δe46-54 was clearly significantly different, with P<0.005 in all cases (we also required an effect size as judged by Cohen’s d > 2). Conversely, no differences were seen between identical edits in the two expression contexts, D16:22 vs D16:24.

#### Global Folding Structure by Protease Challenge

The presence of unfolded regions at low temperatures (i.e. and so do not unfold with increasing temperature, but rather simply contribute to some constant baseline) is not detected by thermal denaturation technique. Such regions are however suggested by differences in helical content. To confirm the presence of these possibly unfolded regions, protease challenge was conducted since proteolytic susceptibility is correlated with flexible and disordered backbone regions. Once again, Δe46-54 is the most well folded with PK50 values (the protease K [PK] giving 50% degradation under our standard assay condition) of 13 ng in D16:22 and 5.1 ng in D16:24 (Figure 4). This may be compared to PK50 values of unskipped 2-STR rods in this region, which range from 8 ng to 22 ng (D16:17 7.6 ng, D17:18 8.2 ng, D20:21 16 ng, D21:22 21 ng, D23:24 8.2 ng), suggesting that this motif is as well folded as unedited rods. The slightly lower value in the D16:24 context may arise from the D22:D23 junction which is not present in the D16:22 series. Mapping studies showed an apparent non-cooperative junction between these two STRs (Mirza et al., 2010). H owever, Δe47-55 and Δe45-53 both display significantly reduced PK50 values in both contexts, in the 0.6 ng to 2.3 ng range. These values are all very different than Δe46-54, and well below these unedited rod PK50 values. This is consistent with significant destabilization of these two edits.

**Figure 4.**
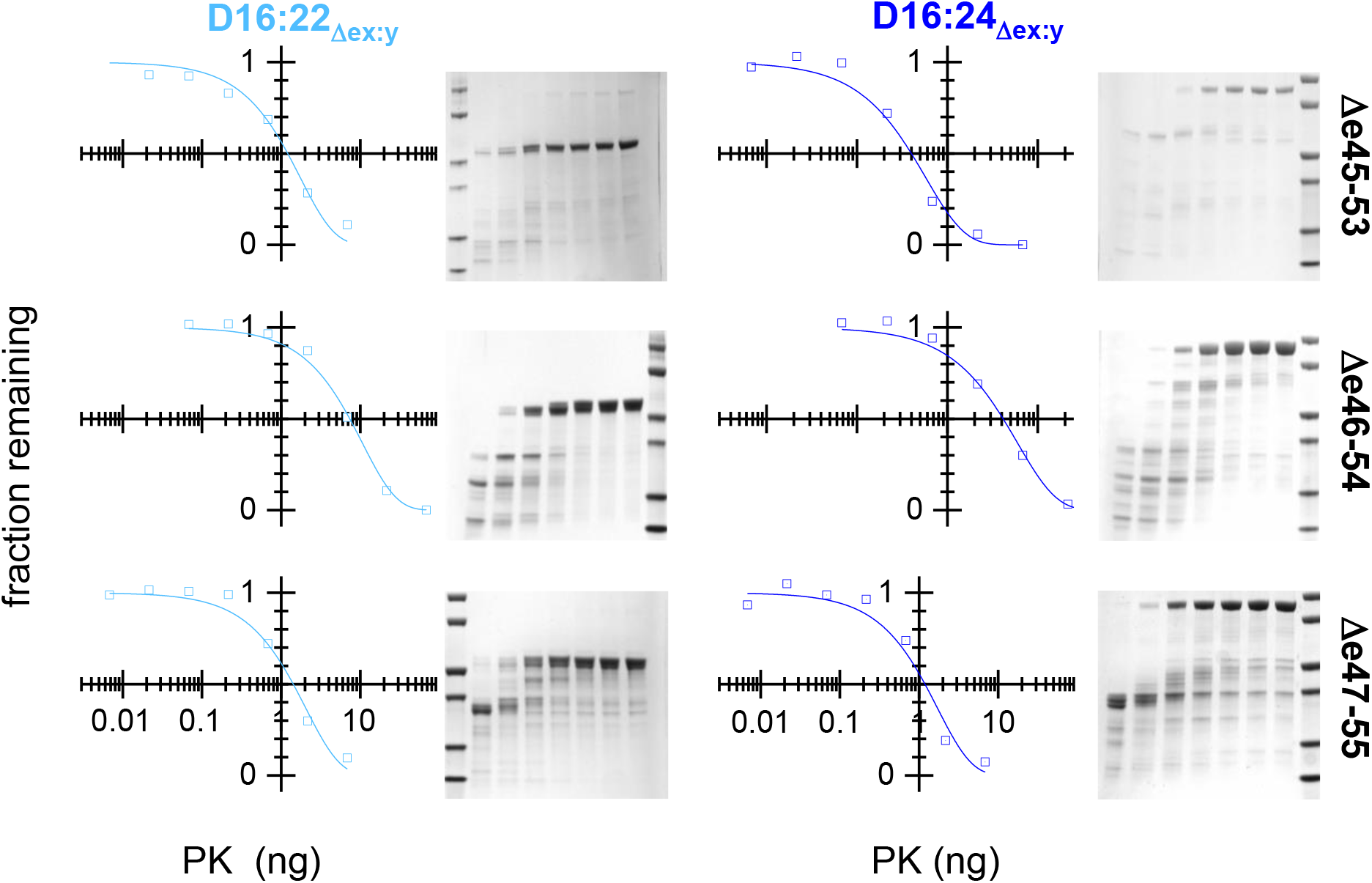
Protease challenge. Target protein were exposed to various concentration of proteinase K under standard condition and analyzed by SDS PAGE. Molecular mass standards (leftmost or right most lane) are at 120, 80, 60, 40, 30, and 20 KDa. In the rest of the gel lanes, proteinase K varies from high to low moving from left to right, as shown on the graphs. The amount of full length protein remaining was assessed by densitometry and fit as described to an exponential decay, yielding a half-life value, PK_50_.

Various measures of stability and well-foldedness are shown together in Table 2, and are quite concordant as shown in Figure 5. In all cases, Δe46-56 is seen to be more stable and well folded than the Δe45-53 and Δe47-55 regardless of whether it was expressed in a D16:22 or D16:24 context. Once again, we judged the statistical relevance by a t-test, with thresholds of P<0.005, or in some cases as noted in Figure 5 or Table 2, P<0.05. In comparing the less stable edits, Δe45-53 was in general less stable than Δe47-55, with the exception of ΔH of unfolding between these two in both parents, and PK50 in the D16:22 context, for which no statistically significant conclusion could be drawn. However, pairwise comparison of Δe45-53 with Δe47-55 is less relevant clinically, since patients will never have a choice between these two, only between one of these and Δe46-54. This strong agreement between such disparate techniques, as well as the agreement between parent molecules gives us confidence that the differences measured reflect underlying differences in the biophysical consequences of these edits on the dystrophin rod.

**Figure 5.**
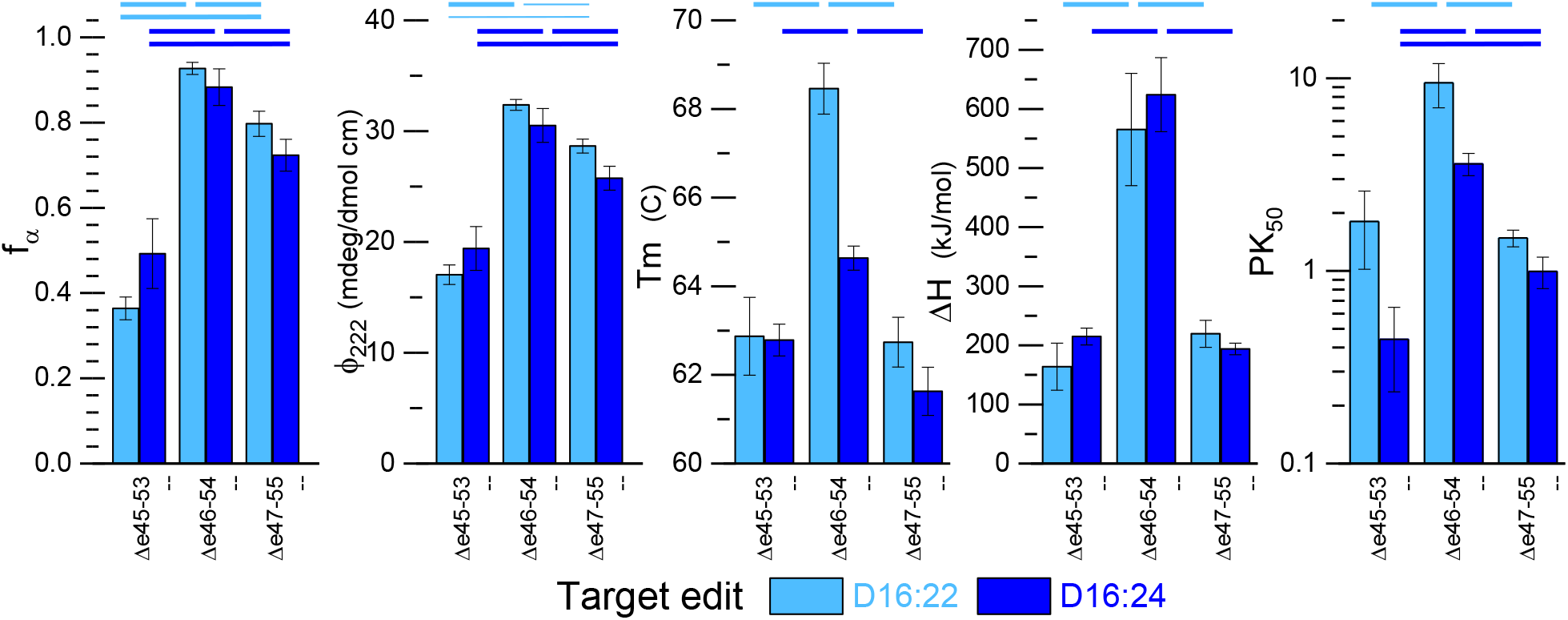
Concordance of Experimental Stability Measures. Various measures of being well structures and stable are compared for the three target edits in both the D16:22 (cyan) and D16:24 (blue) context. In all case, the Δe46-54 edit show greater stability and structure than either Δe45-53 or Δe47-55. All data shown is the mean of n>3 experiments from at least 2 independently produced batches of protein, and error bars reflect 1 standard deviation. Thick lines above indicate statistically significant differences at P<0.005 (two tailed) while thin lines P<0.05, and all with effect size d>2. Complete P and d values are given in supplemental Table S1.

**Table 2.** Experimental Measures of Stability and Well-foldedness. Errors ranges are given as one standard deviation.

| *parent : edit* | $f_{\alpha}$ | $\phi_{222}$<br>(mdeg/dmol cm<sup>2</sup>) | $\Delta H$<br>(kJ/mol) | T<sub>m</sub><br>(°C) | PK<sub>50</sub><br>(ug) |
| --- | --- | --- | --- | --- | --- |
| *D16:22: Δe45-53* | 0.36 ± 0.03 | 17.0 ± 0.9 | 164 ± 40 | 62.9 ± 0.9 | 1.8 ± 0.8 |
| *D16:22: Δe46-54* | 0.93 ± 0.01 | 32.4 ± 0.5 | 565 ± 95 | 68.5 ± 0.6 | 9.5 ± 2.4 |
| *D16:22: Δe47-55* | 0.80 ± 0.03 | 28.7 ± 0.6 | 220 ± 23 | 62.7 ± 0.6 | 1.5 ± 0.1 |
| *D16:24: Δe45-53* | 0.49 ± 0.08 | 19.4 ± 2.0 | 215 ± 14 | 62.8 ± 0.4 | 0.4 ± 0.2 |
| *D16:24: Δe45-54* | 0.88 ± 0.04 | 30.5 ± 1.5 | 624 ± 63 | 64.6 ± 0.3 | 3.6 ± 0.5 |
| *D16:24: Δe45-55* | 0.72 ± 0.04 | 25.8 ± 1.1 | 194 ± 10 | 61.6 ± 0.5 | 1.0 ± 0.2 |

### Computational

#### Initial Models and Helicity

The models produced by Robetta were all molecules that were triple helical STR-like structures (Figure 6). STR proteins are dominated by their three-bundled α-helices, so the simplest metric we examined both experimentally and computationally was total helicity. The experimental data showed significant differences in total helical content as assessed by CD, and our models reproduced the experimental rank order of Δe46-54 > Δe47-55 > Δe45-53, with helicities of 82.8, 81.7 and 80.7 respectively. However, while this order matches (see Figure S3), and is in general agreement for Δe46-54 and Δe47-55, these computational helicity differences are not statistically significant (in contrast to the experimental differences, see Figure 5). Furthermore, the Δe45-53 value is well above the very low experimental value of 36% seen in the D16:22 context, which as previously noted, is not compatible with a typical STR structure. In homology modeling in general, output structures are based on known, well-folded, stable (i.e. unedited) proteins, therefore it is not surprising that we obtained such well folded structures in all cases. Modelling methods thus may bias partially disordered proteins toward their most structured conformer, even while the dynamic behavior of these proteins may result in a lower degree of structure when assessed in a time averaged fashion.

**Figure 6.**
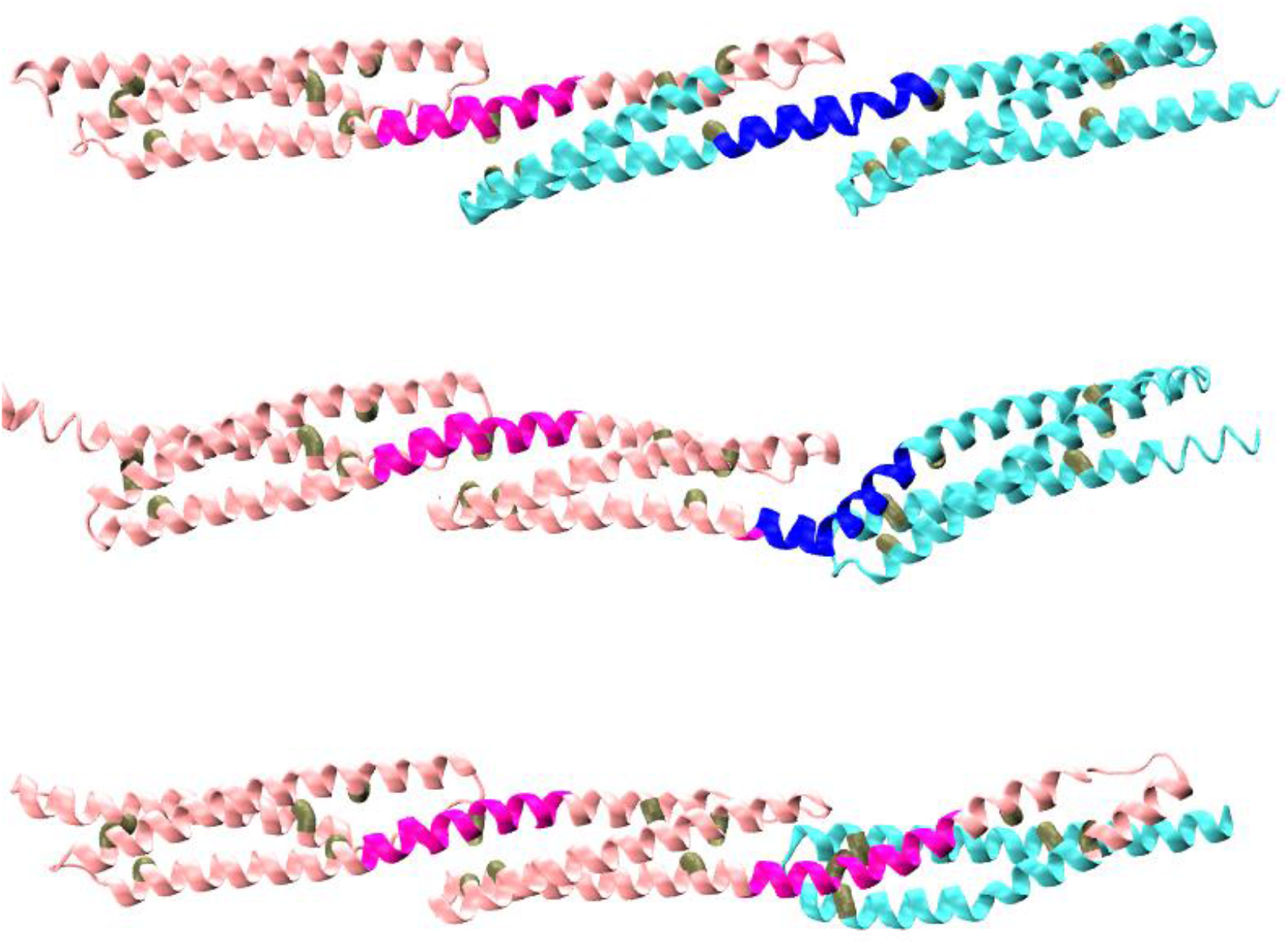
Initial Models. Shown are Δe45-53, (top) Δe46-54 (middle) and Δe47-55 (bottom). The location of the edit site at the transition from pink to blue. The junction sites J1 (left) and J2 (right) are shown in darker pink or blue. The triads of internal hydrophobic sites (see also Figure S2) selected to define each “STR vector” are shown in brown.

For this reason, we then moved to assess the behavior of these models in MD simulations. Here we observed some loss of helicity. This was marginal in terms of the overall helicity, but an examination of this on a residue by residue basis showed that certain regions of the molecule exhibited a high degree of local unfolding, Figure 7. This primarily occurred in the junction regions between the STRs, which is where the third helix of the leading STR propagated into the first helix of the next STR. In the homology model structures, this junction consists of a continuous α-helix that propagates from the third helix of one STR propagates directly into the first helix of the subsequent without a break. This long continuous inter-STR helix is also seen in many crystallographic structures of multi-STR rods from homologous proteins (Djinović-Carugo et al., 1999; Kusunoki et al., 2004; Ortega et al., 2016); and this linkage is thought to contribute to the thermodynamic cooperativity seen in some, but not all, tandem STRs. It is also thought from prior computational studies that this local unwinding of the helix at the junction is an initial event in force-related extension of STR rods, related to their mechanical stabilization role (Mirijanian and Voth, 2008; Paramore et al., 2006).

**Figure 7.**
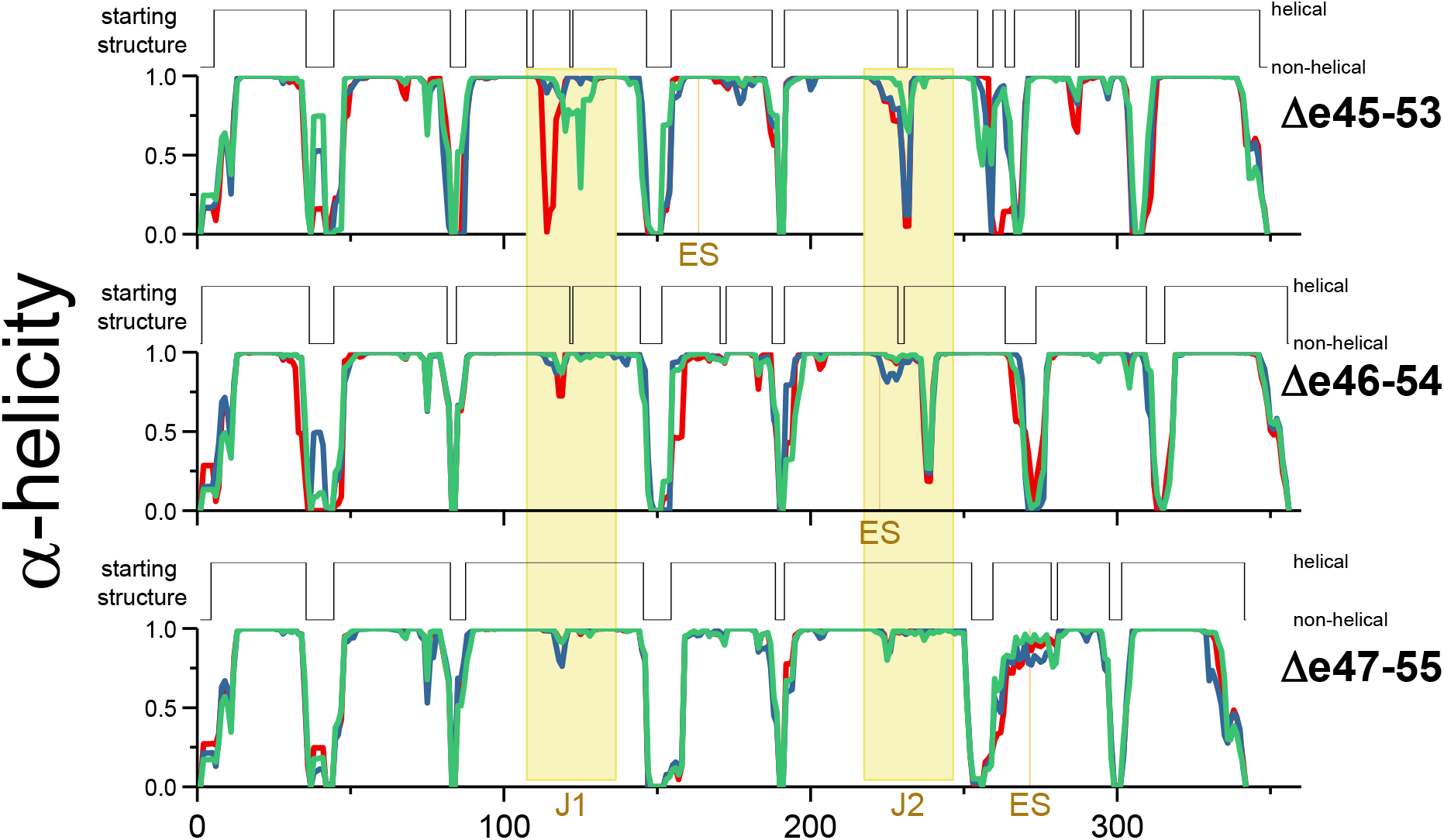
Initial and Dynamic Helicity during MD runs. Shown are the average helicity values at each residue during the three long 1.75 μsec MD runs (red, green and blue plots) for each target protein, compared to the initial model starting structure of each model. The edit site, ES and Junction regions are indicated in yellow.

Since the edit site is also of interest, and it might be reasonably proposed to result in some sort of structural scar in the edited proteins, we also assessed helical unfolding in a 10-residue window centered on the edit sites (five flanking residues on either side). The full distribution of unwinding is shown in the histograms of Figure 8, and is summarized as the mean gap size (how many residues unfold, on average) and fraction of frames with a gap (how often does it unfold) in Table 3.

**Figure 8.**
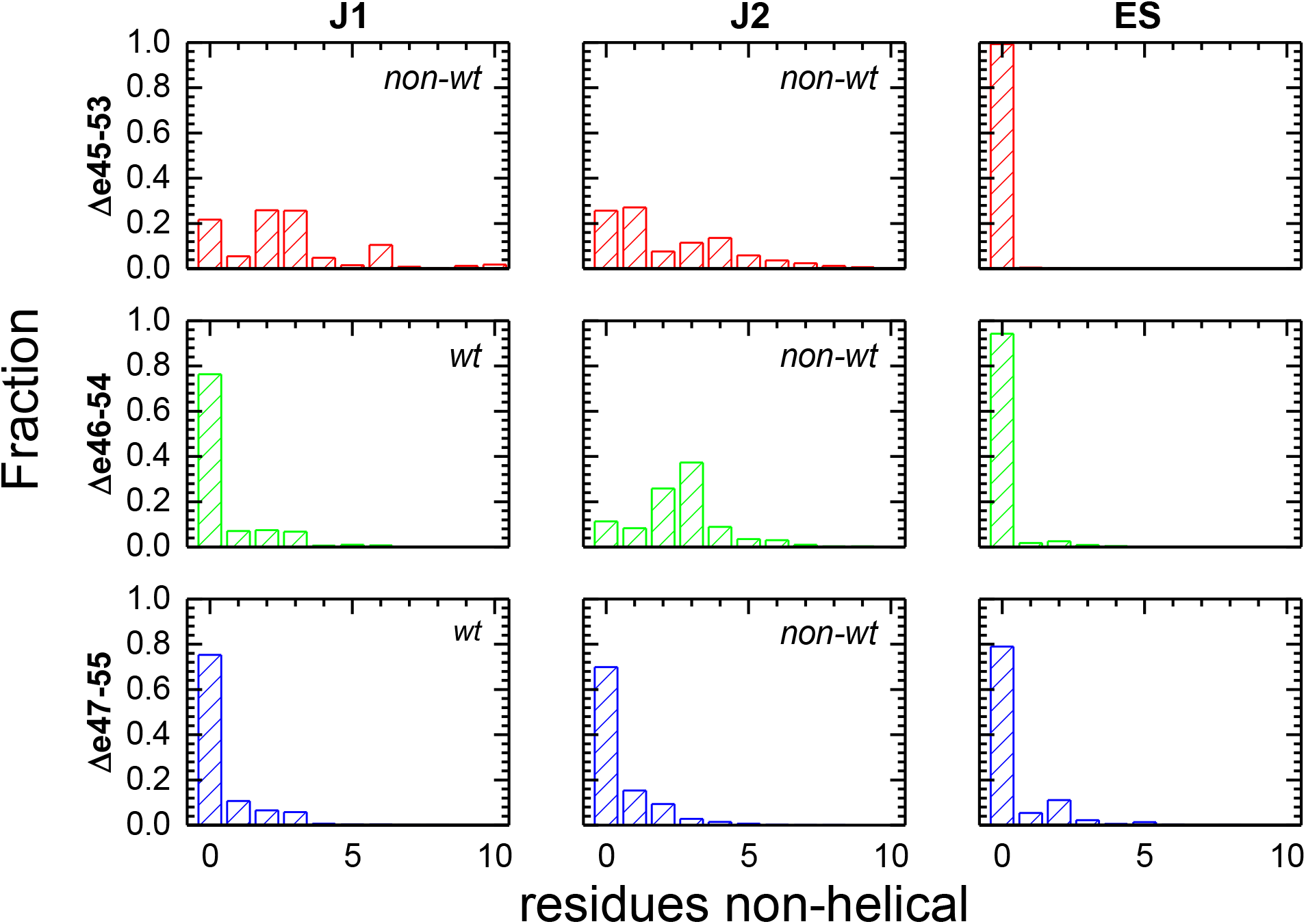
Helicity of edit site, ES and junction regions J1 and J2. The fraction of frames where a gap of non-helical amino acids of a various sizes in the J1, J1 and ES regions is shown All the three edit sites (ES) are largely helical and well-formed, whereas Δe45-53 has both J1 and J2 junctions with significant unfolding – in fact typically more than a full turn (3.6 residues) of an α-helix. This illustrates that a large effect of these edits is felt not locally, at ES, but distal to the specific edit site at the junctions. For Δe46-54 only the J2 junction, which partially overlaps with ES, was significant perturbed. However, the low degree of ES unfolding shows this unfolding is skewed to the junction, not the edit site. Finally, Δe46-54 had junction sites and an edit site that remained largely helical. Junctions are classified as wild-type (wt) or nonwild-type (non-wt) as described in the text, and the non-wt junctions show larger helicity loss.

**Table 3.** Junction and edit site unfolding metrics. Non-wild-type junctions, as described in the text and Figure 8 are highlighted by shading.

|  | mean nonhelical gap size |  |  | fraction of frames with any nonhelical gap |  |  |
| --- | --- | --- | --- | --- | --- | --- |
|  | ES | J1 | J2 | ES | J1 | J2 |
| $\Delta e45-53$ | 0.01 | 2.22 | 1.77 | 0.01 | 0.74 | 0.78 |
| $\Delta e46-54$ | 0.12 | 0.59 | 2.59 | 0.06 | 0.26 | 0.89 |
| $\Delta e47-55$ | 0.46 | 0.50 | 0.55 | 0.21 | 0.25 | 0.30 |

The junction regions exhibited extensive dynamic loss of helicity, especially the least empirically stable edit, Δe45-53. For this edit, the average gap was ~2 residues for both Junction 1 (J1) and Junction 2 (J2), with larger gaps not uncommon. Furthermore, this was a dominant structural feature with ~75 % of frames having a gap. This unfolding was quite heterogeneous and occurred at various spots in this junction region (see supplemental Figure S4 which shows a heatmap of this distribution), with many residues adopting either a helical or unfolded conformation at different times; see for instance the differences between the three runs in the dynamic helicity of J1, and to a lesser extent J2, of Δe45-53 in Figure 7. In contrast, the most empirically stable edit, Δe46-54, displayed a very well-structured J1, with a mean gap size of 0.5 and only 35 % of frames showing any break at all. However, the situation was very different for J2 in this protein: a mean gap of 2.6 residues was present on a more or less permanent basis (90 % of the time). For Δe47-55 (of intermediate stability empirically) we observed well-structured J1 and J2 regions with low (< 0.5 mean gap sizes) and only a minority of frames showing any break at all.

In contrast, the edit site (ES) was comparatively well structured in all cases with < 0.5 residue non-helical, and only infrequently exhibited any non-helical structure. In fact, the least stable protein empirically, Δe45-53, showed almost no ES perturbation (1 % of frames, mean gap 0.01 residue). This strongly suggests that, at least for some exon skip events, the impact of the edit on protein structure does not occur locally. In the other direction, our most stable target, Δe46-54, showed a greater degree of ES helicity loss, at 6% of frames and a mean gap of 0.2 residues; but this is still a comparatively minor perturbation and well lower than the helicity loss at the junction sites. Only for Δe47-55 was significant perturbation seen, where 21 % of frames had some gap, and a mean gap size of 0.46. While this does not approach the size of the gap seen for the perturbed junctions (either junction of Δe45-53 or J2 of Δe46-54), it is comparable to the relatively unperturbed junctions of this protein, and is a larger disturbance than seen for non-edit regions in the three helices of each STR.

We can interpret this junction unfolding as a perturbation caused when edits juxtapose different regions of adjacent STRs at their junctions. Each STR consists of 3 helices, often termed A, B and C; and their connecting loops are often termed the AB-loop and the BC-loop. At each junction, the AB loop of the first STR interacts with the junction spanning helix, as does the BC-loop of the following STR. In some cases, the two loops also interact directly. Looking back at the initial models, Figure 6, we see that for J1 of Δe46-54 and Δe47-55 these (AB-loop, J1 helix, BC-loop) lie on the same side of the edit. This shows that J1 in these cases is the same as it would be in wild-type unedited dystrophin, and all contacts are wild-type. However, for J1 of Δe45-53, the BC-loop comes from the region after the edit, which creates a non-wild-type, non-native interaction. We can thus classify junctions as either **wild type or non-wild-type**. A similar look at J2 shows that all J2’s juxtapose region from both sides of the edit site, and thus are all non-wildtype. In Table 3 we see that wild-type junctions (J1 of Δe 46-54 and Δe47-55) are strongly helical, with low frequency of gap formation and in small gap sizes when formed. In contrast, the non-wild type junctions (J1 of Δe45-53 and J2 of all targets) are less helical with both more frequent breaks and larger gap sizes. This demonstrates that the impact of edits may occur well away from the edit site.

#### Bending

We then moved to understand the impact that this loss of junction structure had on the overall shape of the rod. These rods are thought to play a structural role, linking and stabilizing various cellular components during muscle cell function. As such, their rigidity and shape directly impact their function. We reasoned that a continuous inter STR helix would enforce the classic linear STR rod seen in many crystal structures of non-dystrophin STRs (there are no known multi STR atomic resolution structures from dystrophin, only one single STR dystrophin structure (Muthu et al., 2012)), whereas loss of helicity to a more random coil type junction might allow for bending motion. This may result in an entropic spring type action (Heinrich et al., 2001); or could be a prelude to further unfolding under stress, in a more classical Hookian spring mechanism. To assess such bending, we defined an “STR vector” utilizing specific hydrophobic residues of the amphipathic AbcDefg heptad (see Figures 6 brown residues, and S2 heptad) chosen to locate the two ends of the STR hydrophobic core, and measured the angles between adjacent STR vectors.

Figure 9 illustrates that loss of junction helicity is indeed associated with increased rod flexibility, and further that the least stable edit, Δe45-53, is much more flexible as a result of this unfolding. Looking at the angle correlation plot, where Angle 1 (A1) and Angle 2 (A2) are plotted against each other, we see that for all edits there is a preferred conformer with modest inter STR angles in the 40°-60° range. However, for Δe45-53 we see that there is a distinct second population with a highly bent A1 > 90°. When correlated with junction helicity loss we see that this highly bent A1 conformer only occurs with significant J1 unfolding of > 4 residues (but not J2) (Figure 9, A1 vs J1 map). Loss of helicity is thus seen to allow, but not create, a highly bent structure. For Δe46-54 we do not see a distinct highly bent population, but do observe a long tail of rare conformers with highly bent A1 which are also correlated with J1 helicity loss. For Δe47-55 we see almost no (< 0.1 %) conformers with high bends of > 90°, in keeping with the very low degree of unfolding in this edit. This illustrates that edit-induced junction perturbation impacts the global properties of the rod such as its rigidity.

**Figure 9.**
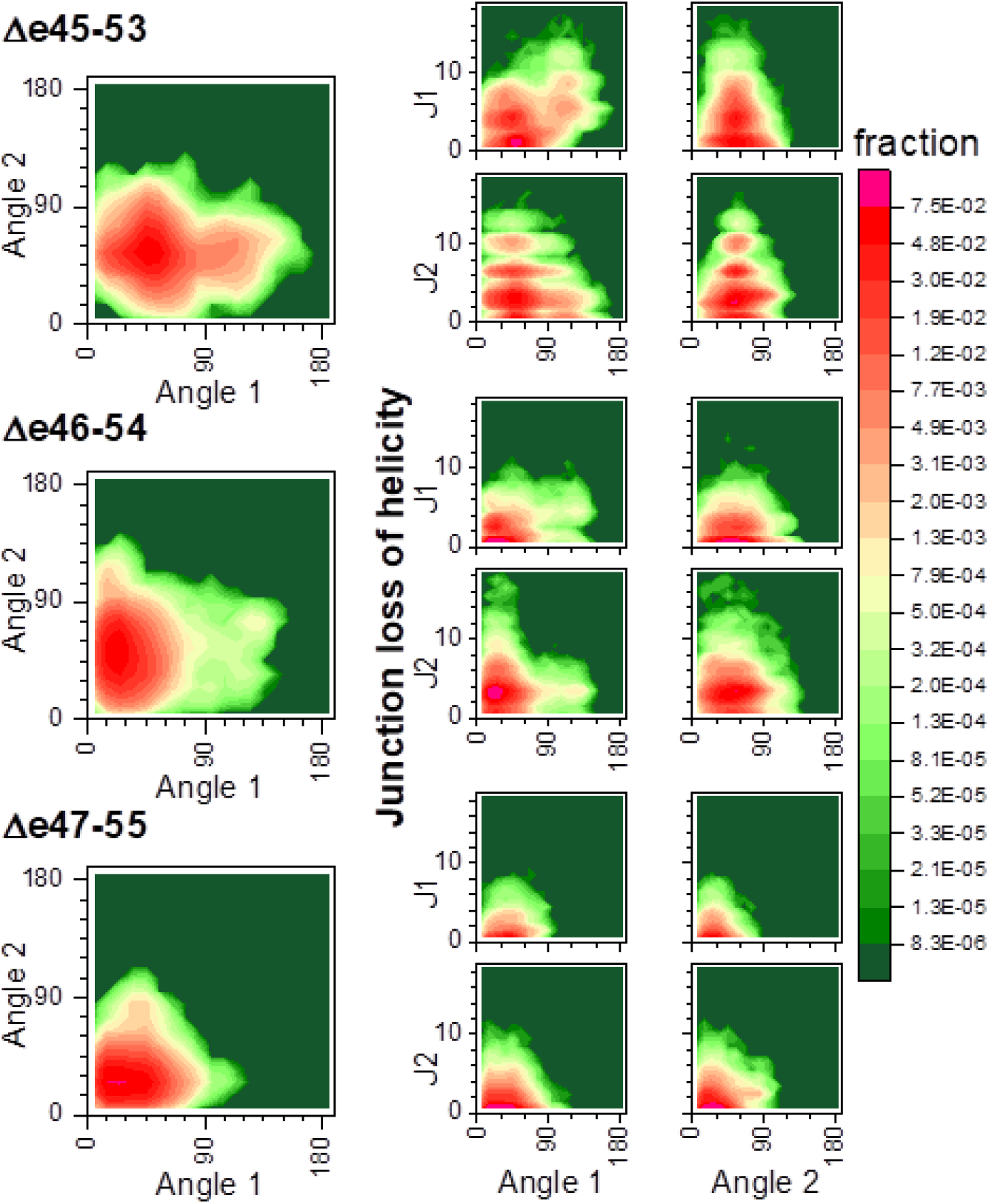
STR bending in relation to junction helicity. The left side is shown correlation between A1 and A2 within each edit, to the right correlation between angle bending and J1 or J2 helicity loss. Δe43-55 show two distinct populations based upon bending of angle A2. We note that in increases in A1 are correlated with increased unfolding at J1 (but not J2), suggesting that loss of helicity here is contributing to increased flexibility of this edit. For A2, both Δe45-53 and Δe46-54 have a broad distribution centered around ~45 degrees but extending beyond 90; these also both the edits that exhibit significant J2 unwinding (Figure 8). In contrast Δe47-55 showed almost no J2 helicity loss (<30 % of frames with any unfolding, and a mean gap a size 0.55; see Table 3) and shown angle 2 distributed in a lower and more narrow range centered around 15 degrees.

So far, for two targets, Δe45-53 (our least empirically stable) and Δe46-54, (most stable) MD has provided a reasonable, if somewhat surprising, narrative for how these edits impact structure and stability: they do not act directly on the edit site, but rather work distally by destabilizing the STR junctions. This leads to junction unfolding and presumably instability. Unfortunately, Δe47-55 does not conform to this explanation. Its junction regions show the least loss of helicity of any target, in both time and extent, and its ES region shows the greatest perturbation – albeit modest compared to junction regions. This perturbation is seen as a dip in the helicity half of the 2^nd^ helix of STR 3 of Δe47-55 (~residue 260-residue 280) in the per residue helicity plot (Figure 7: a much smaller dip is seen near ES of Δe46-54). For this edit, MD shows that the experimental destabilization occurs directly due to ES site perturbation.

#### Motional Correlation Analysis

In two cases, “hybrid STRs” made up of parts of two different naturally occurring STRs occur: Δe45-53 where the second STR is made up of a portion of D17 and portion of D21; and Δe47-55 where the third STR is made up of D18 and D22; see Figures 2 and 6. In order to assess how this might impact the internal STR structure, we turned to motional correlation analysis, Figure 10. Well-formed and tightly interacting domains are expected to exhibit a high degree of correlation, whereas looser or disturbed domains less correlation. Interactions between adjacent secondary structural elements can be seen, and the triple α-helical bundle STR structure provides a unique signature, with three well-defined helix bundling interaction seen: the two antiparallel AB and CB linkages, and the parallel AC linkage producing a characteristic “STR box” signature (see supplemental Figure S5 for a guide to interpreting STR correlation maps). This is easily recognized in most of the correlation maps. In particular those of Δe45-53 and Δe46-54, indicating a high degree of interaction and tight helix-helix interactions.

**Figure 10.**
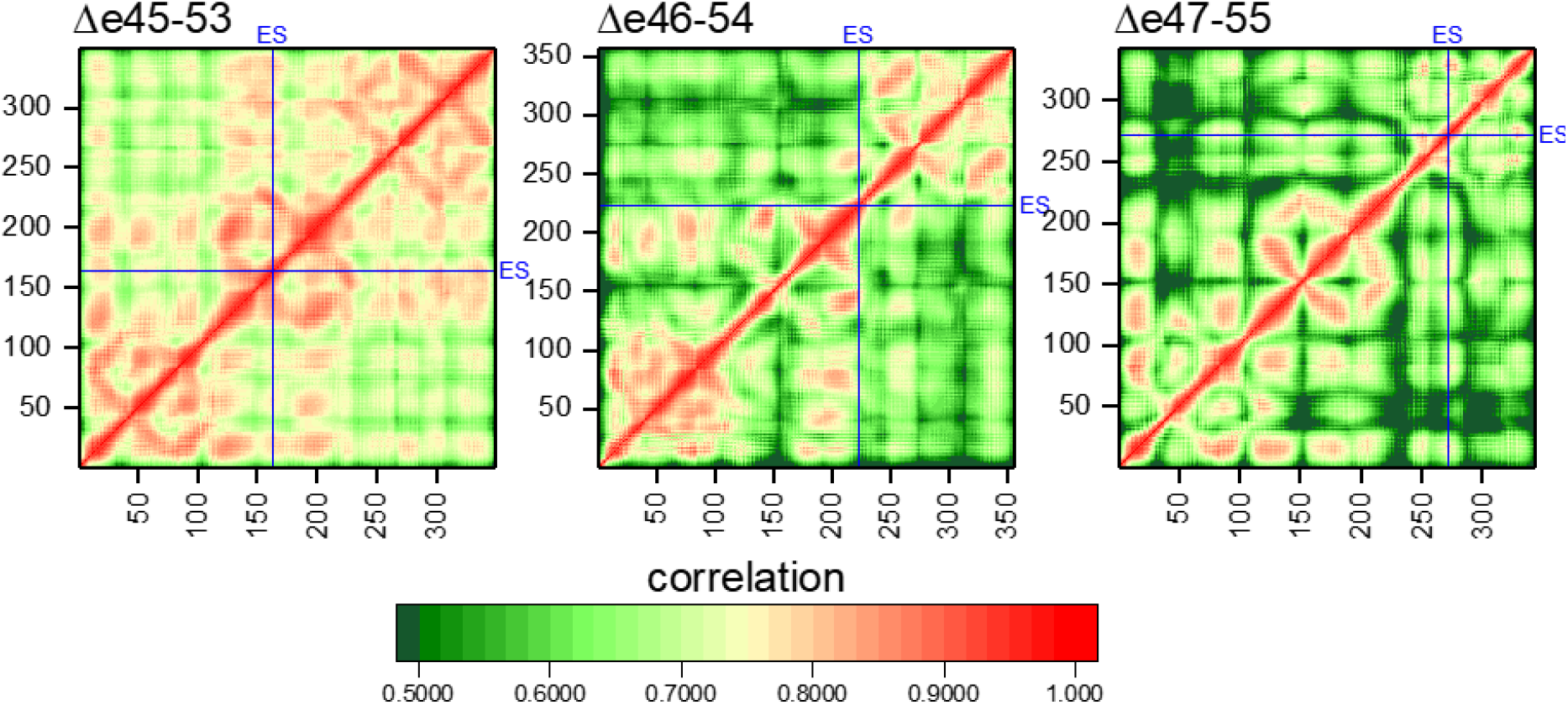
Correlation Analysis of Residue Motion for each Edit. Locations of ES are highlighted with blue lines. Intra-STR Interactions are seen by a characteristic STR box motif (see Figure S5 for interpretation). We see this very easily in Δe45-53 and Δe46-54, and even in the 2^nd^ and somewhat in the 1^st^ STR of Δe47-55. However, the 3^rd^ STR of Δe47-55 is significantly disrupted, with long helical correlation breaking up into much shorter-range interactions. We note that this is not a general consequence of edit site (ES) location, as ES in Δe45-53 is squarely in the middle of STR2, which is not perturbed. Off-diagonal inter-STR interactions are seen strongly for J1, but are absent for J2 of Δe46-54, in agreement with the increased unfolding of J2 but not J1 for this edit (as seen in Figure 8).

In contrast, for Δe47-55 we see this box signature only in the first and second STRs, while the third, which contains the ES in this target, is highly disrupted. Rather than three long individual, and highly correlated, regions, we see a scattering of smaller, local regions and less long-range motional correlation in all three helices, indicating a loss of long range internal interactions. The Δe45-53 target also contains a hybrid STR with an internal ES, but in this case it long range correlations are retained, and this edit does not appear to disrupt global STR interactions. In the case of Δe46-54, the edit site lies in the second STR but relatively close to the STR2-STR3 junction and appears not to perturb long range interactions either.

We also note that the overall level of correlation is higher for Δe45-53 than for the other proteins, while Δe47-55 is the lowest. This is simply a result of the increased flexibility of Δe45-53 (and decreased flexibility of Δe47-55) as demonstrated by, and corroborating, the STR angle analysis. This results in correlated motion of all atoms in each STR as they bend relative to the rod as a whole. This is confirmed as this excess correlation disappears when the correlation calculated only within individual STRs (see Supplemental Figure S6).

#### Energetics of STR motifs

We also quantified the internal energetics of each STR using MMGBSA energy analysis of each STR in each target protein. This is not a true measure of stability of folding, which is the difference in energy between the folded and unfolded states, as we do not at this time have computational access to the unfolded state. However, the total internal energy can give some indication of the forces involved in maintaining the STR structure and how the edits may perturb them in certain cases. Here, we see that all molecules have similar internal energies for their first STR (Figure 11A). This is sensible, since the first STR in all cases is a wild-type STR, the 16^th^ in native dystrophin and has an identical amino acid sequence in all targets. However, we note that the second STR has a significantly lower energy in Δe45-53; and the third STR has lower stabilizing energy in Δe47-55. These two are precisely the STRs where the edit site is located, thus are nonwild-type, hybrid STRs made up of sections of two different wild-type STRs: a hybrid D17/21 in the case of Δe45-53; a hybrid D18/22 in Δe47-55. This shows that while these hybrid STRs can form, they seem to have reduced stability, the extent of which may vary depending on the specific nature of the STRs hybridized. This corroborates the loss of internal STR structure seen in the correlation matrices for these STRs. Once again, only Δe46-54 escapes this, with none of its three STRs energetically perturbed. In that case, the edit occurs near the end of D18, which is largely intact, with only a comparatively small section of D21 (~5 amino acids) fused to it. In this sense, STR 2 of Δe46-54 is “almost” wild-type.

**Figure 11.**
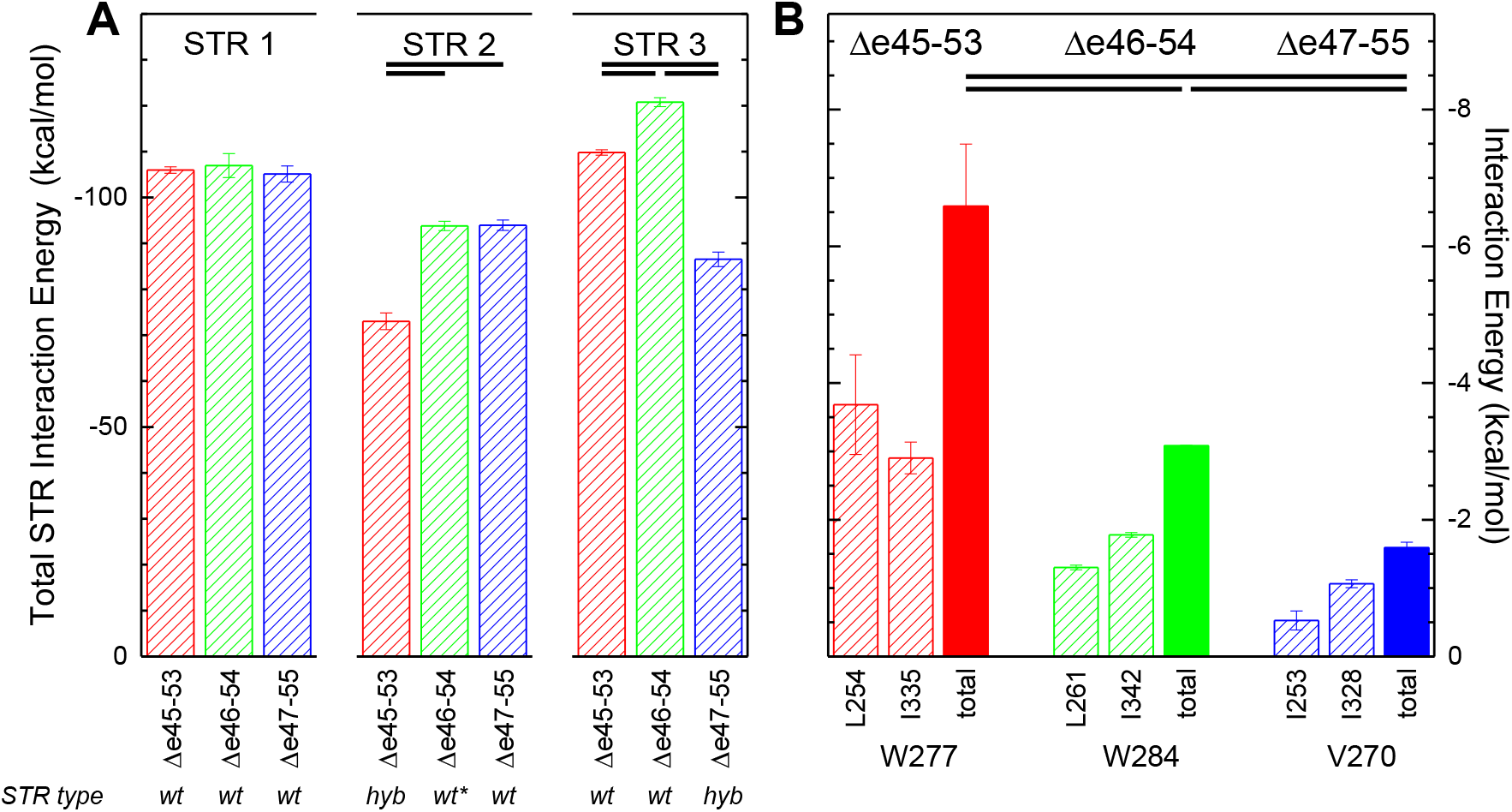
MMGBSA Energy of Interaction. **A** Total Pairwise interaction for all residues within each STR of each edit. Values shown are averages calculated over the three 1.75 usec runs, with error bars of one standard deviation. Statistical significance at the criteria specified (P<0.005, d> 2; all values in Table S2) is indicated by bars near the top. STR 1 is a wild-type (wt) STR and does not contain any edit sites, and has the same primary sequence for all targets – and the total STR energies are quite similar. The edit site in Δe45-53 occurs in STR 2, which is a hybrid (hyb) STR in this case, being made up of part of wild type D17 and D21; it exhibits lower energy than STR2 of Δe46-54 or Δe47-55. Similarly, STR3 of Δe47-55 is a hybrid STR in this target; it too exhibits lower energy than STR 3 of the other targets. For Δe46-54 the edit site occurs near the end of STR 2, very close to its junction STR 3 and STR 2 is very nearly wild-type, indicated as wt*. **B** Individual pairwise interaction energies for the W277 and W284 heptad triads of Δe45-53 and Δe46-54, and the homologous V270 triad of Δe47-55 (see Figure S2 for the position of these in the alignment, and Figures 6 for a view of these positions in the models). We note that when the edit replaced this W with a smaller and less hydrophobic V, the interaction energy is reduced, and is an example of how the overall destabilization occurs for these edited STRs.

We also were able to identify specific amino acid level interactions that are perturbed in the Δe47-55 case. We noticed that the region near the C-terminus of STR3 – the perturbed, hybrid STR in this target – was particularly destabilized, and appeared to partially unbundle, up to the first hydrophobic triad (see Figure S7). Here we noted that this site is a tryptophan in the other edits which, but has been substituted by a much smaller and less hydrophobic valine in Δe47-55. Tryptophan residues are particularly well conserved in STR motifs, and have been proposed to strongly contribute to their stability (Pantazatos and MacDonald, 1997). We thus examined the energetic interactions between these triad partners and found that they were significantly reduced, Figure 11B. While the reduction is only a modest fraction of the reduction seen in the overall STR energy, it shows how perturbation of these heptad triads can be significant.

#### Explicit Solvent Simulations

We conducted most of our experiments using an implicit solvent model, as a trade-off between physical reality and computational feasibility. Implicit solvent models are clearly less physically realistic and may skew results due to solvation effect; however explicit solvent models suffer from slower simulation speeds and therefore lower sampling, thereby introducing a different source of error. To understand whether solvent effect biases impact our results, we conducted additional explicit solvent simulations starting from various conformers derived from cluster analysis of the implicit solvent standard dataset. If any of these were non-physical – or more accurately, if any of these conformers were the result of differences between the implicit and explicit solvent force field – then we would expect that those conformers would move away from their initial cluster when subject to explicit MD simulations. This did not happen, as shown by the root mean square deviation (RMSD) analysis in Table 4. All runs stayed within their starting cluster on average, with the sole exception of the third cluster of Δe46-54 which was a minor conformer representing only 2.2 % of the implicit solvent dataset. An evolution of this RMSD distance over time is shown in Supplemental Figure S8.

**Table 4.** Explicit MD Movement Explicit MD runs were begun from conformation points produced by cluster analysis of the implicit standard data set. Clusters with > 1.5% abundance were used ( 4 for Δe45-53 and Δe46-52 and 3 for Δe47-55). The abundance and size (rmsd) of each cluster is indicated. We then monitored the average rmsd of each trajectory from the representative structures of each of the other clusters. If solvation differences between implicit and explicit runs perturbed the energetics, then some clusters would be expected to rapidly evolve away from their starting points toward another cluster. The lowest rmsd (closest) cluster is highlighted in bold. In fact, most stayed closest to their starting structure, and within the size of the that starting cluster. The sole exceptions are the lowest abundance cluster in De45-53, which appeared to evolve to a point midway between cluster 1 and cluster 4; and the third cluster of De46-54 which appears to have evolved into another very low abundance (2%) cluster 4. This suggests that solvation model differences effects are not extreme.

| cluster | abundance | size | rmsd from |  |  |  |
| --- | --- | --- | --- | --- | --- | --- |
|  |  |  | 1 | 2 | 3 | 4 |
| $\Delta e_{45-53}$ | | | | | | |
| 1 | 0.62 | 21.1 | **13.5** | 27.0 | 21.3 | 23.2 |
| 2 | 0.17 | 19.7 | 30.6 | **20.9** | 26.2 | 35.8 |
| 3 | 0.06 | 18.0 | 23.5 | 24.3 | **17.2** | 29.9 |
| 4 | 0.06 | 19.6 | **20.5** | 32.5 | 27.6 | **20.5** |
| $\Delta e_{46-54}$ | | | | | | |
| 1 | 0.88 | 20.6 | **13.9** | 24.7 | 19.8 | 22.6 |
| 2 | 0.03 | 21.4 | 25.8 | **16.1** | 28.4 | 31.7 |
| 3 | 0.02 | 18.7 | 22.1 | 29.4 | 25.0 | **14.8** |
| 4 | 0.02 | 19.4 | 21.3 | 28.7 | 24.5 | **14.3** |
| $\Delta e_{47-55}$ | | | | | | |
| 1 | 0.92 | 19.1 | **12.0** | 19.6 | 20.0 |  |
| 2 | 0.03 | 18.8 | 18.3 | **17.9** | 24.7 |  |
| 3 | 0.02 | 20.4 | 18.4 | 25.1 | **16.0** |  |

The fact that none of these conformers was rapidly perturbed (i.e. within the three 250s ns simulations per system) when moved to an explicit solvent model suggests that the differences between these solvation models are not severe, and do not greatly impact the conformers obtained. While all conformers are in equilibrium and interconvertible during runs – either explicit or implicit – the longer more rapid pace of implicit runs allowed us to more fully sample the conformational space. We also examined the conformations of all edits during the explicit runs, redoing the helicity and bending analysis, and no great differences were observed. This is shown in dynamic per residue helicity, Figure S9, homologous to Figure 6; junction and edit site helicity loss, Figure S10, homologous to Figure 8; and in the overall rod bending Figure S11, homologous to Figure 9.

## Discussion

Our results clearly show that Δe46-54 is more stable and structurally more similar to unskipped dystrophin rods than the other two edits studied, with Δe47-55 being intermediate and Δe45-53 being the least stable and least well-formed. Consistent results were obtained across several empirical stability measures, and regardless of the parent rod in which it was expressed, D16:22 or D16:24. Helicity, PK50, and thermal denaturation measurements, which probe the order, disorder, and stabilization energy of the folded form, give the same overall order, Δe46-54 > Δe47-55> Δe45-53. This consistency between the parent rod and measurement types gives us confidence these differences are intrinsic to the edit, and not some peculiarity of the context in which the edit is studied, or the techniques used to probe it.

Taken together we get the following empirical picture of these edits:

Δe45-53 is an edit that produces a perturbed structure of low helicity inconsistent with STR structure, has significant disorder and significantly reduced stabilization enthalpy.

Δe46-54 is an edit that produced a typical STR type structure that is well formed with few disordered regions, high helicity and typical unfolding thermodynamics.

Δe47-55 is also a perturbative edit resulting in a significant disorder, but while the structure is of reduced stability when thermally challenged, it does have appropriate helicity at low temperatures, and so might be considered to be a marginal edit.

### MD to Understand the Molecular Origins of Experimental Effects

However, as much as we can determine the phenomenological properties of these edits by this type of characterization, we cannot understand at a molecular level why some edits seem to produce viable structures – or conversely why some do not. Unfortunately, there are no high-resolution structures of dystrophin rods in this region to help understand how such edits might manifest at an atomic level, although there are many known homologous STR structures from many other proteins. To work around this, we developed homology-based models and used molecular dynamics simulations to help understand the molecular basis of these differences.

The models produced were consistent with STR structures seen in other proteins and were grossly consistent with the experimental evidence that these are α-helix dominated structures. However, they were more helical than our experimentally produced structures as assessed by CD. We believe this is partially due to a bias in many homology algorithms which may favor highly structured outputs. This idea is supported by our MD simulations, which showed that helicity dropped somewhat once the molecules were equilibrated to 300 K and further reduced by local unfolding in a dynamic fashion during long runs. While we did come closer to the helicity values for the least empirically structured motifs, we did not get full agreement. However, we were only able to simulate on the order of 2 μs, which is very short compared to the time scale of the *in vitro* experiments (minutes to hours).

Nonetheless, our computational results did reproduce the stability order seen experimentally and have the virtue of being able to provide insights as to why and how this instability arises in some edits but not others. The most important conclusion is that in some cases, the exon editing has impacts well away from the edit site. In our least stable motif, Δe45-53, the edit site was in fact very well structured. This edit site occurs in the middle of the second STR in the target and is almost completely helical even in dynamic fashion during runs. However, the edit induced changes in the STR junction regions. By juxtaposing new residues not normally in contact with each other, these junctions are destabilized and begin to unfold, and MD runs show high degree of loss of helicity at both junctions for this edit.

However, this is not always the case. The most stable edit, Δe46-54 has only modest perturbations. For the edit of intermediate experimental stability, Δe47-55, the junctions do not seem perturbed at all, whereas the edit site in this case does seem to experience some loss of structure. Correlation analysis demonstrated loss of long range interaction, and energy analysis showed a reduced overall inter-residue interaction energy within the edit containing STR. This paints a complex picture – in some cases the edits act locally, whereas in others they are locally accommodated in STR structure, but their effects propagate to STR junctions where they have a large impact on structure.

### Relevance to Exon Skipping Therapy

As exon skipping therapy has been developed, most of the effort has gone into simply getting it to work: finding a pharmacologically acceptable compound that is effective in inducing some exon skipping in enough tissue to provide a clinically meaningful benefit. With the recent approval of eteplirsen a milestone has been reached, although there is certainly a long way to go in improving both delivery and skipping efficiency, as well as in expanding the range of exon targeted to bring this treatment option to more patients. We are now at the stage where skipping for many, if not all, exons is scientifically within reach (Aartsma-Rus et al., 2017; Nakamura, 2017; Niks and Aartsma-Rus, 2017). However even in the limit of perfect exon skipping, the protein that is created will have some edit and may therefore have functional differences from wild type dystrophin. This results in DMD patients essentially becoming therapeutically created BMD patients – and in many cases, patients and clinicians may have some choice in determining which form of BMD to aspire to.

Retrospective studies of BMD patients provide some guidance on how to choose, but also demonstrate great heterogeneity and uncertainty. There are many thousands of known mutations for DMD and BMD, meaning that many specific edits are rare. Given these small sample sizes and the fact that patients with the same underlying exon deletion can progress at highly different rates (Findlay et al., 2015a; Kaspar et al., 2009a; Nicolas et al., 2015) it is challenging in most cases to compare one edit against another. To get around this sampling problem, the various BMD defects have typically been analyzed in structurally related groups to increase sample size and statistical power. This has demonstrated correlations with the underlying structure of the protein on a general level (Kaspar et al., 2009a). However, this grouping may obscure the underlying physical causes of an ailment if dystrophin molecules with diverse tertiary structures are analyzed together.

Clinical relevance has been most clearly demonstrated in a relationship between BMD defects and dilated cardiomyopathy (Kaspar et al., 2009b) (DCM), a leading cause of mortality in BMD. In that study, to achieve statistical power many edits were grouped based upon their features in relation to the STR structure of the rod, and certain groups were associated with a large delay in onset of DCM. “In-phase” edits that were expected to have less perturbed STR structures progressed more rapidly, with an 11-year difference in age of DCM diagnosis than “out-of-phase” edits that were expected to have more perturbed STR structures. While this clearly showed a link between DCM severity and edit structure, this study also emphasized that patients with the same underlying defect can progress at very different rates. For instance, the most rapidly progressing patient in the more favorable in-phase cohort (patient 109; DCM at 15 years of age, Figure 3A in that paper) was more severely afflicted than any of the supposedly worse off out-of-phase group. However, in aggregate, this clinical evidence shows that the underlying protein structure is important (i.e. the significant delay in DCM), even if variability of progression may be an inherent part of BMD.

We believe that factors relating to structural characteristics of the edit at the protein level are what underly this clinically significant correlation. Dystrophin is a structural protein whose primary functions is to provide a mechanical linkage in muscle cell. It also has multiple scaffolding and organizational functions e.g. organizing the DGC and NOS (Greenberg et al., 1994; Judge et al., 2011; Lai et al., 2009). All of these rely on proper folding, structure and stability of this protein and might easily be perturbed in edited proteins, either natural, as in BMD; or therapeutically produced, as in exon skipping or other therapies.

### Relation to Clinically Observed Cases

We note that two of the edits corresponding to the ones studied here have been clinically observed: Δe45-53 and Δe46-54 and are found 95 and 4 times, respectively, in the public Leiden database ((White and den Dunnen, 2006); Sept 2017 update). Δe47-55 is not found. The underlying DMD-type deletions which might be repaired by exon skipping to these edits have also been clinically observed: Δe46-53 is noted 25 times, and Δe47-54 is found 5 times. Of course, this database is not an unbiased survey of the human population, but is composed of only clinically observed cases, and so is biased toward those seeking medical attention. All DMD patients require medical attention, and many BMD patients do as well, as they progress. However, it is an open question as to whether there is a cohort of subclinical BMD-type deletions out there who are unknown to the scientific community since they are so subclinical, they do not seek help. Such subclinical edits are the ‘holy grail’ of exon skipping, but might be expected to be underrepresented in this database.

The creation frequency of deletions is related to two factors: size of the regions in which the breakpoints must lie (i.e. the size of the flanking introns), and the length of the deletions, with the length dependence scaling with an exponent of −1.5 (Zhang and Gerstein, 2003). Using this, we might thus calculate a cross-section for each exon edit from the known sizes of dystrophin exons and introns; this is shown in Table 5. Other factors such as chromatin structure and CpG islands are also important, but appear to only make a minor contribution to mutational hotspots (Kondrashov, 2003). This explains the observed prevalence somewhat, as we can see that intron 46i is quite small, reducing the observed prevalence of Δe47-54 and Δe47-55; whereas intron 44i is quite large, increasing the prevalence of Δe45-53.

**Table 5.** Observed frequencies of relevant edits and breakpoint intron sizes. Cross-section IS calculated form intron sizes and overall length as described in the text; and prevalence is as observed in the Leiden clinical database at the time of writing as described in the text.

| Deletion | 5' intron |  | 3' intron |  | Length |  | Cross-section | Prevalence | Ratio |
| --- | --- | --- | --- | --- | --- | --- | --- | --- | --- |
|  | Size |  | Size |  | Max | Min |  |  |  |
| ***DMD*** |  |  |  |  |  |  |  |  |  |
| $\Delta e46-53$ | 45i | 36111 | 53i | 21230 | 310195 | 252852 | 5.1 | 25 | 4.9 |
| $\Delta e47-54$ | 46i | 2334 | 54i | 30127 | 304218 | 271755 | 0.5 | 5 | 11.0 |
| ***BMD*** |  |  |  |  |  |  |  |  |  |
| $\Delta e45-53$ | 44i | 248401 | 53i | 21230 | 558772 | 289139 | 19.1 | 95 | 5.0 |
| $\Delta e46-54$ | 45i | 36111 | 54i | 30127 | 340477 | 274237 | 6.4 | 4 | 0.6 |
| $\Delta e47-55$ | 46i | 2334 | 55i | 120219 | 424627 | 302072 | 1.3 | 0 | 0.0 |

Selection is important as well. The DMD alleles are under extremely strong X-linked selection, since virtually all males with DMD do not reproduce. As expected, this 100 % culling of males results in a mutation-selection equilibrium in which one third of cases are *de novo* mutations, with the rest being created in female carriers in the prior few generations and little long term inheritance (GRIMM et al., 2012). However, for BMD-type edits this is not the case, with a much greater carrier incidence observed, ~ 90 %, (Lee et al., 2014) demonstrating that BMD alleles overall are under lower selection pressure, which should elevate their incidence relative to DMD. When examining the ratio of database incidence to this cross section, we find the opposite: the DMD type deletions, and one of our BMD-type edits are represented at a similar frequency, and two edits in particular seem underrepresented: Δe46-54 and Δe47-55. The evidence for abnormally low prevalence of Δe47-55 is weak, since none are reported whereas only ~ 6 might be expected using a prevalence to cross-section ratio of 5. However, for Δe46-54 the evidence is stronger: 4 cases have been reported whereas 32 might have been expected. While this is not conclusive, it does suggest that this Δe46-54 defect has not come in contact with medical system in proportion to its expected incidence, and supports our conclusions based on the biophysical evidence that it is a is mild edit with low phenotypic consequences. Conversely, the fact that Δe45-53 is observed in the same ratio to its expected frequency as DMD defects in the same region suggests that it is severe enough that these patients seek medical attention at similar rates to severely affected DMD patients. We have identified one report of this edit in the literature that has clinically meaningful progression data. In that case, the Δe45-53 patient has an age of DCM onset of 36 years (Kaspar et al., 2009a), significantly below the median of 43 years in the relevant cohort in that study, making it a comparatively severe case as expected from the above analysis.

### Consequences

Empirically, this study shows that alternative exon skip repairs can result in repaired proteins of very different stabilities. In this case, we found that the Δe46-54 repair is much more stable than either Δe45-53 or Δe47-55. This means that DMD defects Δe46-53 or Δe47-54 can be repaired in two different ways, with different consequences with respect to the nature of the protein produced. While the experimental evidence is clear on which edit is more stable, it does not shed light on the molecular details of why this is. Computational studies suggest that this can arise in different ways. Perturbation of the edit site is the most obvious way, and is the case for Δe47-55, where most of the disrupted residues are in the ES region. Juxtaposing amino acids that did not evolve to be in proximity might quite naturally be expected to produce suboptimal interactions and reduced stability. However, a subtler effect is seen in Δe45-53, which has the largest experimentally observed destabilization. In this case it the edit produces changes in the global tertiary structure that destabilize regions distal to the edit site, in particular the junction regions between adjacent STRs. In all known high-resolution structures of multi-STRs rods, the last helix of the leading STR propagates directly into the first helix of the next. Furthermore, the loop regions of the other helices (AB loop of the leading STR, BC loop of the following STR) are in close proximity to and interact with these junction regions. This then provides a model for how these edits perturb these structures: by placing different and unnatural combinations of these two loops and the junction helix together at each junction, essential stabilizing interactions that are present in naturally-evolved tandem STR junctions are lost. In some cases, this may fortuitously result in productive interactions (i.e. in our most stable edit, Δe46-54) whereas in others it might be destabilizing. This shows the idiosyncratic nature of exon editing, and the values of both experimental and computational analysis of each edit to identify such compatible and incompatible junctions. Overall, our MD results suggest that the consequences of exon edit can impact structure in a number of distinct ways.

This heterogeneity may be less than satisfying in terms of providing a simple or uniform explanation for exon edit stability, and suggests that a simple heuristic understanding of exon edit impact at the protein level may not be possible. However, it does illuminate possible origins of these differences, and that each repair can be understood on a case by case basis. This highlights the need for a thorough analysis of each edit, and suggests that a combined experimental and computational approach can provide meaningful comparisons where alternative exon skipping repair is possible.

This is most crucial in cases where a clinical choice may be made. Currently there is only one choice for exon skipping, eteplirsen (Exondys51) which targets exon 51. However clinical trials for compounds targeting exons 44, 45 and 53 are underway (i.e. NCT02329769, NCT02500381, NCT01957059), and preclinical programs exist for many other exons. If development of the therapies targeting these new exons is successful (and there is no reason to think they will pose any greater regulatory hurdles than exon 51) the prospect of patients and their physicians facing a choice about which to use will become a reality. Since the disease is slowly progressive, it seems likely that the consequences of this choice will only manifest after years of therapy. Some serious symptomology such as DCM generally only develops in the second or later decades of life, and may be exacerbated – or ameliorated – by choice of therapy made decades earlier, and careful consideration of this choice may be important for optimal clinical outcomes.

## Acknowledgments

The computational aspects of this work were supported by the National Institute of General Medical Sciences of the National Institutes of Health (grant 1R35GM119647). The content is solely the responsibility of the authors and does not necessarily represent the official views of the National Institutes of Health. This work used the Extreme Science and Engineering Discovery Environment, which is supported by National Science Foundation Grant No. ACI-1053575.

**Author Contributions**
The manuscript was written through contributions of KM JW and NM. JW led the computational side, NM led the experimental side. KM and ET conducted the experimental work, KM conducted the computational work. All authors have given approval to the final version of the manuscript.

## References

Aartsma-Rus, A., and Krieg, A.M. (2017). FDA Approves Eteplirsen for Duchenne Muscular Dystrophy: The Next Chapter in the Eteplirsen Saga. Nucleic Acid Ther. 27, 1–3.

Aartsma-Rus, A., Straub, V., Hemmings, R., Haas, M., Schlosser-Weber, G., Stoyanova-Beninska, V., Mercuri, E., Muntoni, F., Sepodes, B., Vroom, E., et al. (2017). Development of Exon Skipping Therapies for Duchenne Muscular Dystrophy: A Critical Review and a Perspective on the Outstanding Issues. Nucleic Acid Ther. 27, 251–259.

Anthony, K., Arechavala-Gomeza, V., Ricotti, V., Torelli, S., Feng, L., Janghra, N., Tasca, G., Guglieri, M., Barresi, R., Armaroli, A., et al. (2014). Biochemical Characterization of Patients With In-Frame or Out-of-Frame DMD Deletions Pertinent to Exon 44 or 45 Skipping. JAMA Neurol. 71, 32–40.

Birnkrant, D.J., Bushby, K., Bann, C.M., Apkon, S.D., Blackwell, A., Brumbaugh, D., Case, L.E., Clemens, P.R., Hadjiyannakis, S., Pandya, S., et al. (2018). Diagnosis and management of Duchenne muscular dystrophy, part 1: diagnosis, and neuromuscular, rehabilitation, endocrine, and gastrointestinal and nutritional management. Lancet Neurol. 17, 251–267.

Calvert, R., Kahana, E., and Gratzer, W.B. (1996). Stability of the dystrophin rod domain fold: evidence for nested repeating units. Biophys. J. 71, 1605–1610.

Case, D.A., Cheatham, T.E., Darden, T., Gohlke, H., Luo, R., Merz, K.M., Onufriev, A., Simmerling, C., Wang, B., and Woods, R.J. (2005). The Amber biomolecular simulation programs. J. Comput. Chem. 26, 1668–1688.

Chowdhury, R., Rasheed, M., Keidel, D., Moussalem, M., Olson, A., Sanner, M., and Bajaj, C. (2013). Protein-protein docking with F(2)Dock 2.0 and GB-rerank. PloS One 8, e51307.

Djinović-Carugo, K., Young, P., Gautel, M., and Saraste, M. (1999). Structure of the alpha-actinin rod: molecular basis for cross-linking of actin filaments. Cell 98, 537–546.

Djinovic-Carugo, K., Gautel, M., Ylänne, J., and Young, P. (2002). The spectrin repeat: a structural platform for cytoskeletal protein assemblies. FEBS Lett. 513, 119–123.

Falzarano, M.S., Scotton, C., Passarelli, C., and Ferlini, A. (2015). Duchenne Muscular Dystrophy: From Diagnosis to Therapy. Molecules 20, 18168–18184.

Findlay, A.R., Wein, N., Kaminoh, Y., Taylor, L.E., Dunn, D.M., Mendell, J.R., King, W.M., Pestronk, A., Florence, J.M., Mathews, K.D., et al. (2015a). Clinical phenotypes as predictors of the outcome of skipping around DMD exon 45. Ann. Neurol. 77, 668–674.

Findlay, A.R., Wein, N., Kaminoh, Y., Taylor, L.E., Dunn, D.M., Mendell, J.R., King, W.M., Pestronk, A., Florence, J.M., Mathews, K.D., et al. (2015b). Clinical phenotypes as predictors of the outcome of skipping around DMD exon 45. Ann. Neurol. 77, 668–674.

Fontana, A., Polverino de Laureto, P., De Filippis, V., Scaramella, E., and Zambonin, M. (1997). Probing the partly folded states of proteins by limited proteolysis. Fold. Des. 2, R17–26.

Gibson, D.G., Glass, J.I., Lartigue, C., Noskov, V.N., Chuang, R.-Y., Algire, M.A., Benders, G.A., Montague, M.G., Ma, L., Moodie, M.M., et al. (2010). Creation of a bacterial cell controlled by a chemically synthesized genome. Science 329, 52–56.

Götz, A.W., Williamson, M.J., Xu, D., Poole, D., Le Grand, S., and Walker, R.C. (2012). Routine Microsecond Molecular Dynamics Simulations with AMBER on GPUs. 1. Generalized Born. J. Chem. Theory Comput. 8, 1542–1555.

Greenberg, D.S., Sunada, Y., Campbell, K.P., Yaffe, D., and Nudel, U. (1994). Exogenous Dp71 restores the levels of dystrophin associated proteins but does not alleviate muscle damage in mdx mice. Nat. Genet. 8, 340–344.

Greenfield, N., and Fasman, G.D. (1969). Computed circular dichroism spectra for the evaluation of protein conformation. Biochemistry (Mosc.) 8, 4108–4116.

Grimm, T., Kress, W., Meng, G., and Müller, C.R. (2012). Risk assessment and genetic counseling in families with Duchenne muscular dystrophy. Acta Myol. 31, 179–183.

Hamuro, L., Chan, P., Tirucherai, G., and AbuTarif, M. (2017). Developing a Natural History Progression Model for Duchenne Muscular Dystrophy Using the Six-Minute Walk Test. CPT Pharmacomet. Syst. Pharmacol. 6, 596–603.

Harper, S.Q., Hauser, M.A., DelloRusso, C., Duan, D., Crawford, R.W., Phelps, S.F., Harper, H.A., Robinson, A.S., Engelhardt, J.F., Brooks, S.V., et al. (2002). Modular flexibility of dystrophin: implications for gene therapy of Duchenne muscular dystrophy. Nat. Med. 8, 253–261.

Heinrich, V., Ritchie, K., Mohandas, N., and Evans, E. (2001). Elastic thickness compressibilty of the red cell membrane. Biophys. J. 81, 1452–1463.

Hubbard, S.J. (1998). The structural aspects of limited proteolysis of native proteins. Biochim. Biophys. Acta 1382, 191–206.

Humphrey, W., Dalke, A., and Schulten, K. (1996). VMD: visual molecular dynamics. J. Mol. Graph. 14, 33–38, 27–28.

Hunter, J.D. (2007). Matplotlib: A 2D Graphics Environment. Comput. Sci. Eng. 9, 90–95.

Ipsaro, J.J., Huang, L., Gutierrez, L., and MacDonald, R.I. (2008). Molecular epitopes of the ankyrin-spectrin interaction. Biochemistry (Mosc.) 47, 7452–7464.

John, D.M., and Weeks, K.M. (2000). van’t Hoff enthalpies without baselines. Protein Sci. Publ. Protein Soc. 9, 1416–1419.

Judge, L.M., Arnett, A.L.H., Banks, G.B., and Chamberlain, J.S. (2011). Expression of the dystrophin isoform Dp116 preserves functional muscle mass and extends lifespan without preventing dystrophy in severely dystrophic mice. Hum. Mol. Genet. 20, 4978–4990.

Kabsch, W., and Sander, C. (1983). Dictionary of protein secondary structure: pattern recognition of hydrogen-bonded and geometrical features. Biopolymers 22, 2577–2637.

Kaspar, R.W., Allen, H.D., Ray, W.C., Alvarez, C.E., Kissel, J.T., Pestronk, A., Weiss, R.B., Flanigan, K.M., Mendell, J.R., and Montanaro, F. (2009a). Analysis of dystrophin deletion mutations predicts age of cardiomyopathy onset in becker muscular dystrophy. Circ. Cardiovasc. Genet. 2, 544–551.

Kaspar, R.W., Allen, H.D., and Montanaro, F. (2009b). Current understanding and management of dilated cardiomyopathy in Duchenne and Becker muscular dystrophy. J. Am. Acad. Nurse Pract. 27, 241–249.

Khajavi, M., Inoue, K., and Lupski, J.R. (2006). Nonsense-mediated mRNA decay modulates clinical outcome of genetic disease. Eur. J. Hum. Genet. 14, 1074.

Kim, D.E., Chivian, D., and Baker, D. (2004). Protein structure prediction and analysis using the Robetta server. Nucleic Acids Res. 32, W526–W531.

Komaki, H., Nagata, T., Saito, T., Masuda, S., Takeshita, E., Sasaki, M., Tachimori, H., Nakamura, H., Aoki, Y., and Takeda, S. ‘ichi (2018). Systemic administration of the antisense oligonucleotide NS-065/NCNP-01 for skipping of exon 53 in patients with Duchenne muscular dystrophy. Sci. Transl. Med. 10.

Kondrashov, A.S. (2003). Direct estimates of human per nucleotide mutation rates at 20 loci causing mendelian diseases. Hum. Mutat. 27, 12–27.

Kusunoki, H., Minasov, G., Macdonald, R.I., and Mondragón, A. (2004). Independent movement, dimerization and stability of tandem repeats of chicken brain alpha-spectrin. J. Mol. Biol. 344, 495–511.

Lai, Y., Thomas, G.D., Yue, Y., Yang, H.T., Li, D., Long, C., Judge, L., Bostick, B., Chamberlain, J.S., Terjung, R.L., et al. (2009). Dystrophins carrying spectrin-like repeats 16 and 17 anchor nNOS to the sarcolemma and enhance exercise performance in a mouse model of muscular dystrophy. J. Clin. Invest. 119, 624–635.

Lange, O.F., and Grubmüller, H. (2006). Generalized correlation for biomolecular dynamics. Proteins Struct. Funct. Bioinforma. 62, 1053–1061.

Le, S., Yu, M., Hovan, L., Zhao, Z., Ervasti, J., and Yan, J. (2018). Dystrophin As a Molecular Shock Absorber. ACS Nano 12, 12140–12148.

Le Rumeur, E. (2015). Dystrophin and the two related genetic diseases, Duchenne and Becker muscular dystrophies. Bosn. J. Basic Med. Sci. 15, 14–20.

Lee, T., Takeshima, Y., Kusunoki, N., Awano, H., Yagi, M., Matsuo, M., and Iijima, K. (2014). Differences in carrier frequency between mothers of Duchenne and Becker muscular dystrophy patients. J. Hum. Genet. 59, 46–50.

Linde, L., and Kerem, B. (2008). Introducing sense into nonsense in treatments of human genetic diseases. Trends Genet. TIG 24, 552–563.

Maier, J.A., Martinez, C., Kasavajhala, K., Wickstrom, L., Hauser, K.E., and Simmerling, C. (2015). ff14SB: Improving the Accuracy of Protein Side Chain and Backbone Parameters from ff99SB. J. Chem. Theory Comput. 11, 3696–3713.

McCourt, J.L., Rhett, K.K., Jaeger, M.A., Belanto, J.J., Talsness, D.M., and Ervasti, J.M. (2015). In vitro stability of therapeutically relevant, internally truncated dystrophins. Skelet. Muscle 5, 13.

Menhart, N. (2006). Hybrid spectrin type repeats produced by exon-skipping in dystrophin. Biochim. Biophys. Acta 1764, 993–999.

Miller, B.R., McGee, T.D., Swails, J.M., Homeyer, N., Gohlke, H., and Roitberg, A.E. (2012). *MMPBSA.py* : An Efficient Program for End-State Free Energy Calculations. J. Chem. Theory Comput. 8, 3314–3321.

Mirijanian, D.T., and Voth, G.A. (2008). Unique elastic properties of the spectrin tetramer as revealed by multiscale coarse-grained modeling. Proc. Natl. Acad. Sci. 105, 1204–1208.

Mirza, A., Sagathevan, M., Sahni, N., Choi, L., and Menhart, N. (2010). A biophysical map of the dystrophin rod. Biochim. Biophys. Acta 1804, 1796–1809.

Mongan, J., Simmerling, C., McCammon, J.A., Case, D.A., and Onufriev, A. (2007). Generalized Born Model with a Simple, Robust Molecular Volume Correction. J. Chem. Theory Comput. 3, 156–169.

Muthu, M., Richardson, K.A., and Sutherland-Smith, A.J. (2012). The crystal structures of dystrophin and utrophin spectrin repeats: implications for domain boundaries. PloS One 7, e40066.

Nakamura, A. (2017). Moving towards successful exon-skipping therapy for Duchenne muscular dystrophy. J. Hum. Genet. 62, 871–876.

Nguyen, H., Roe, D.R., and Simmerling, C. (2013). Improved Generalized Born Solvent Model Parameters for Protein Simulations. J. Chem. Theory Comput. 9, 2020–2034.

Nicolas, A., Delalande, O., Hubert, J.-F., and Le Rumeur, E. (2014). The spectrin family of proteins: a unique coiled-coil fold for various molecular surface properties. J. Struct. Biol. 186, 392–401.

Nicolas, A., Raguénès-Nicol, C., Ben Yaou, R., Ameziane-Le Hir, S., Chéron, A., Vié, V., Claustres, M., Leturcq, F., Delalande, O., Hubert, J.-F., et al. (2015). Becker muscular dystrophy severity is linked to the structure of dystrophin. Hum. Mol. Genet. 24, 1267–1279.

Niks, E.H., and Aartsma-Rus, A. (2017). Exon skipping: a first in class strategy for Duchenne muscular dystrophy. Expert Opin. Biol. Ther. 17, 225–236.

Ortega, E., Manso, J.A., Buey, R.M., Carballido, A.M., Carabias, A., Sonnenberg, A., and de Pereda, J.M. (2016). The Structure of the Plakin Domain of Plectin Reveals an Extended Rod-like Shape. J. Biol. Chem. 291, 18643–18662.

Pantazatos, D.P., and MacDonald, R.I. (1997). Site-directed mutagenesis of either the highly conserved Trp-22 or the moderately conserved Trp-95 to a large, hydrophobic residue reduces the thermodynamic stability of a spectrin repeating unit. J. Biol. Chem. 272, 21052–21059.

Paramore, S., Ayton, G.S., and Voth, G.A. (2006). Extending a Spectrin Repeat Unit. II: Rupture Behavior. Biophys. J. 90, 101–111.

Peck, A. (2008). Beginning GIMP: from novice to professional (Berkeley, Calif. : New York: Apress, Distributed to the Book trade worldwide by Springer-Verlag).

Ray, A., Lindahl, E., and Wallner, B. (2012). Improved model quality assessment using ProQ2. BMC Bioinformatics 13, 224.

Roe, D.R., and Cheatham, T.E. (2013). PTRAJ and CPPTRAJ: Software for Processing and Analysis of Molecular Dynamics Trajectory Data. J. Chem. Theory Comput. 9, 3084–3095.

Sahni, N., Mangat, K., Le Rumeur, E., and Menhart, N. (2012). Exon edited dystrophin rods in the hinge 3 region. Biochim. Biophys. Acta 1824, 1080–1089.

Sarkis, J., Hubert, J.-F., Legrand, B., Robert, E., Chéron, A., Jardin, J., Hitti, E., Le Rumeur, E., and Vié, V. (2011). Spectrin-like repeats 11-15 of human dystrophin show adaptations to a lipidic environment. J. Biol. Chem. 286, 30481–30491.

Shao, J., Tanner, S.W., Thompson, N., and Cheatham, T.E. (2007). Clustering Molecular Dynamics Trajectories: 1. Characterizing the Performance of Different Clustering Algorithms. J. Chem. Theory Comput. 3, 2312–2334.

Shimizu-Motohashi, Y., Miyatake, S., Komaki, H., Takeda, S. ‘ichi, and Aoki, Y. (2016). Recent advances in innovative therapeutic approaches for Duchenne muscular dystrophy: from discovery to clinical trials. Am. J. Transl. Res. 8, 2471–2489.

Towns, J., Cockerill, T., Dahan, M., Foster, I., Gaither, K., Grimshaw, A., Hazlewood, V., Lathrop, S., Lifka, D., Peterson, G.D., et al. (2014). XSEDE: Accelerating Scientific Discovery. Comput. Sci. Eng. 16, 62–74.

White, S.J., and den Dunnen, J.T. (2006). Copy number variation in the genome; the human DMD gene as an example. Cytogenet. Genome Res. 115, 240–246.

Wolny, M., Grzybek, M., Bok, E., Chorzalska, A., Lenoir, M., Czogalla, A., Adamczyk, K., Kolondra, A., Diakowski, W., Overduin, M., et al. (2011). Key amino acid residues of ankyrin-sensitive phosphatidylethanolamine/phosphatidylcholine-lipid binding site of βl-spectrin. PloS One 6, e21538.

Zhang, Z., and Gerstein, M. (2003). Patterns of nucleotide substitution, insertion and deletion in the human genome inferred from pseudogenes. Nucleic Acids Res. 31, 5338–5348.

Zimowski, J.G., Pilch, J., Pawelec, M., Purzycka, J.K., Kubalska, J., Ziora-Jakutowicz, K., Dudzińska, M., and Zaremba, J. (2017). A rare subclinical or mild type of Becker muscular dystrophy caused by a single exon 48 deletion of the dystrophin gene. J. Appl. Genet. 58, 343–347.

