## Supplementary material for "Empirical and Computational Comparison of Alternative Therapeutic Exon Skip Repairs for Duchenne Muscular Dystrophy": Supp Methods Figures and Tables (from another version of this preprint: version 1, 2019-01-23)

### Supplemental Methods

#### Cloning and Protein Production

These methods have been set out in earlier work on other exon edited rods (Sahni et al., 2012). Primers were designed that could PCR amplify the N- and C-terminal regions before or after the edit site, with Gibson overlap regions spanning the exon edit. These fragments amplified by PCR, cloned into a pGEX expression vector by Gibson assembly (Gibson et al., 2010), and transformed into *E. coli* NEB express (a commercially modified DH5α strain optimized for protein expression).

Our methods for purifying these proteins have been previously described in details as applied to both native and exon edited rods (Sahni et al., 2012). Use of double affinity tagging, with a different tag at each end, has been found to be crucial for exon edited proteins, many of which are somewhat unstable and susceptible to degradation *in vivo* while being expressed, or in the early stages (i.e. the lysis) of the purification. Any proteolytic nicking (e.g. at a potentially disrupted edit site) will split it into two separate fragments, each of which has only one tag; only full-length, undegraded protein will be selected by both affinity protocols.

After growth and expression, the bacteria are harvested by centrifugation and lysed by sonication in the presences of a non-ionic detergent, (1% triton X-100) and then the crude lysate clarified by centrifugation and applied to a glutathione agarose chromatography support, washed to remove extraneous proteins, and eluted with 10 mM glutathione. The crude preparation was then treated with thrombin to release the GST affinity tag, and then applied to a Hitrap Co IMAC (immobilized metal affinity chromatography, Thermo scientific) column to bind the second affinity tag, the C-terminal $His_9$ region. This was then washed with 10mM imidazole to remove weakly bound proteins and eluted with a gradient to 500 mM imidazole. It is important to remove the GST tag, since it is large (26 kDa, 226 amino acids) and it would significantly interfere with the signal of the dystrophin protein domains we are interested in, which are all 3 (D16:22 parent series) or 5 (D16:24 parent series) STRs in size, and so ~330 or ~550 amino acids in size. On the other hand, the small 9 amino acid $His_9$ tag was not removed, since this is <3% and only makes a very minor contribution to the signal in the assays used to characterize these targets.

After this, a final step of ion exchange purification (HiTrap Q column, thermo scientific; tris pH 8 buffer, with a gradient to 500 mM NaCl) was used to remove any remaining non-full-length species present, due for instance to alternative minor thrombin sensitivity sites during GST removal or other degradation process that might occur in the time in between the two affinity steps.

#### Implicit Solvent Molecular Dynamics Simulations

Structures provided by Robetta modelling were minimized and equilibrated through a multi-stage protocol:

- Side chains were relaxed while the backbone atoms were restrained by a 10 kcal/(mol $A^2$) restraint for 10,000 steps.
- 1,000 minimization steps where all protein atoms were free.
- The temperature of the system was increased from 5 to 300 K during a 50 ps Langevin dynamics calculation with heavy backbone atoms restrained with a 10 kcal/(mol $A^2$) restraint.
- The backbone restraints were gradually released over 450 ps of further equilibration at 300K.

#### Explicit Solvent Molecular Dynamics Simulations

The starting representative structures from the cluster analysis were rotated to align them along the x axis of a solvent box with a minimum distance of 10 Å from any protein atom to the edge of the box, and then solvated with TIP3P (Jorgensen et al., 1983) water molecules. Additional sodium and chloride ions were added to neutralize and create an 0.15 M NaCl environment. Structures were minimized and equilibrated through a multi-stage protocol similar to the one in implicit solvent mentioned above, except that:

1. an initial minimization of all solvent molecules while the protein atoms were restrained was performed for 10,000 steps
2. and the system was heated to 300 K in the 50 ps NPT ensemble with backbones retrained by a 10 kcal/(mol·$Å^2$) restraint and gradually released to equilibration for 950 ps.

Three independent simulations were then conducted at constant pressure for 250 ns utilizing the SHAKE algorithm and Monte Carlo barostat (Åqvist et al., 2004), with only the last 150 ns used for analysis. A 2 fs timestep, with a hard cutoff of 10.0 Å and long-range electrostatics handled with the particle mesh Ewald-method (Essmann et al., 1995) were used. This resulted in 450 ns of data over the three runs for each cluster.

#### Computational Analysis

##### *Helicity*

Helicity analysis was accomplished by cpptraj, which detects helices by the DSSP criteria (Kabsch and Sander, 1983). Overall helicity for each target over the standard dataset was calculated, as well as the helicity by residue for each run. Stable α-helices were identified as those residues with >80% average helicity over the standard data set.

##### *Junction loss of helicity*

In quantifying this junction loss of helicity, we ran up against the problem that there is no precise definition as to where each STR starts and ends to precisely locate '*the*

*STR junction*'. There are two common definitions in common use: Koenig and Kunkel, KK (Koenig et al., 1988) and Winder Gibson Kendrick-Jones, WGK (Winder et al., 1995) that differ by up to eight amino acids. As such, we opted for an empirical definition, by first locating the three helices of each STR by the point at which that average per residue helicity over the standard data set dropped below 90 % (shown in highlighting in Figure S2). We then defined the junction as the region between the third helix of one STR and the first helix of the next, plus a window of four residues (i.e. one turn) and quantified unfolding as the length of the longest gap of nonhelical residues on a per frame basis (quantified in Figure 8).

*Inter-STR Bending*

We also examined the overall rod bending by quantifying an angle between the individual STRs. To unambiguously determine this angle, we first defined an "STR vector" for each of the three STRs by identifying a common group of core hydrophobic residues for each STR of each target protein in a consistent fashion. The helices of STRs are amphipathic, with a distinctive repeating `AbcDefg` heptad pattern of amino acid polarity where the A and D position are hydrophobic and oriented to the interior of each bundle (Djinovic-Carugo et al., 2002). We identified triads of the interacting of such positions at each end of the STR in a common position, as shown in the empirical sequence alignment in Figure S2, and in the model structures in Figure 6. The vector between the center of mass of the C$\alpha$ atoms of each of these two triads in each STR defined a "STR vector" and the STR bending angles 1 and 2 were defined as the angles between these vectors for adjacent STRs.

*MMGBSA Free Energy.*

To examine the interaction between residues contributing to the energetic stability/instability of each alternative repair, we examined pairwise energies between residues within each STR for each standard data set by the Molecular Mechanics/Generalized Born Surface Area, MMGBSA (Chowdhury et al., 2013). The Generalized Born implicit solvent model using the atomic radii (Mongan et al., 2007) was used for the analysis. The non-polar contribution to solvation was calculated using the linear combination of pairwise overlaps method with a probe radius of 1.4 Å and surface tension of 0.0072 kcal/(mol·Å$^2$). The MMPBSA.py script was used for this analysis (Miller et al., 2012). We calculated the pairwise energetic interactions of all residues with within each STR as defined by the empirical helicity alignment (Figure S2) that arose out of the helicity analysis; this gives an indication of the forces stabilizing each STR. This technique is also able to isolate any particular pairwise interaction between any selected amino acids, which are presented for a few special cases.

*Correlated Motion Analysis.*

Translational and rotational motions were first removed by alignment of alpha carbon atoms to the average structure of the trajectory. This aligned trajectory dataset was then analyzed for general motional correlation (Lange and Grubmüller, 2006) which calculated the mutual information correlation matrix describing how each amino acid residue's motion is correlated with every other. This provides a picture of how different parts of the molecule interact. Correlation was calculated both over the entire molecule, as well as independently for each STR as defined by the empirical alignment in Figure S2.

*Cluster Analysis.*

Cluster analysis was performed for each target's standard data set. Since our helicity and bending analysis revealed the dynamics of the molecule was characterized by changes in the secondary structure of these junction regions, and to a lesser extent, of the edit site regions, we performed cluster analysis on the phi and psi backbone dihedral angles of these resides. Specifically, we used the same 10 amino acid edit site, and a 21 residue junction windows used for helicity analysis above, as shown on Figures 7, S2 and S4. The hierarchical agglomerative algorithm (Shao et al., 2007) with average-linkage was used for the analysis with an average angular distance of 200° between clusters.

### Supplemental Tables

**Table S1**

Statistical parameters for relevance assessment of the empirical stability data shown in Figure 5. Effect size was assessed by Cohen's d parameter $d = \frac{\mu_1 - \mu_2}{s}$ where s is the pooled standard deviation $s = \left(\frac{(n_1-1)s_1+(n_2-1)s_2}{n_1+n_2-2}\right)^{0.5}$. P values were calculated by a two tailed Student's t-test. Values meeting our significance criteria (P<0.05; d>2) are in black, others are in grey. Highly significant P values < 0.005 are also **bolded**. The most relevant pairwise comparison are Δe45-53 to Δe46-54 and Δe45-53 to Δe46-54, since these represent alterative exon edits of some disease state where clinical choice is possible. These are presented in the two comparisons to the left. In the comparison to the right, Δe45-53 is compared to Δe47-55; this is perhaps less interesting since it does not reflect any clinical choice; nonetheless is may be of fundamental scientific interest in helping us understand how certain edits are less stable than others. Non-significant values (by the criteria above) are greyed out.

| | Δe45-53 to Δe46-54 | | Δe47-55 to Δe46-54 | | Δe47-55 to Δe46-54 | | |
|---|---|---|---|---|---|---|---|
| | d | P-value | d | P-value | d | P-value | |
| $f_{\alpha}$ | 28 | **2.9E-07** | 6.0 | **5.3E-04** | 15 | **4.7E-05** | *D16:22* |
| | 6.4 | **4.1E-04** | 3.9 | **3.7E-03** | 3.6 | 1.1E-02 | *D16:24* |
| $\phi_{222}$ | 22.7 | **8.1E-07** | 6.8 | **3.0E-04** | 15 | **5.0E-05** | *D16:22* |
| | 6.4 | **3.8E-04** | 3.5 | 6.1E-03 | 4.0 | 8.1E-03 | *D16:24* |
| $T_m$ | 7.5 | **4.0E-05** | 10 | **7.4E-06** | 0.2 | 8.1E-01 | *D16:22* |
| | 5.6 | **7.1E-04** | 7.0 | **1.0E-03** | 2.6 | 1.9E-02 | *D16:24* |
| ΔH | 5.5 | **2.3E-04** | 5.0 | **4.0E-04** | 1.7 | 5.1E-02 | *D16:22* |
| | 9.9 | **4.8E-05** | 9.6 | **3.0E-04** | 1.7 | 8.1E-02 | *D16:24* |
| $PK_{50}$ | 4.2 | **9.9E-04** | 4.2 | **2.7E-03** | 0.5 | 5.2E-01 | *D16:22* |
| | 8.6 | **4.6E-04** | 7.2 | **9.2E-04** | 2.8 | 2.6E-02 | *D16:24* |

**Table S2**

Statistical assessment of the relevance of MMGBSA energies shown in Figure 11. We use the same relevance criteria (d>2, P<0.005) as explained the text and in table S1. In Figure 11A we presented total pairwise energy over all residues for each STR, and compared individual STRs (STR1, STR2 or STR3) between targets. In Figure11B we examined one particular interaction involving a hydrophobic triad positions normally containing a conserved W that has previous been identified as especially significant in STR folding and stability (see discussion, and Figures 6 and S2), and compared this between targets. Non-significant values (by the criteria above) are greyed out.

| comparison | total pairwise energy over STR (figure 9A) | | | | | | triad total MMGBSA Energy (figure 9B) | |
|---|---|---|---|---|---|---|---|---|
| | STR1 | | STR2 | | STR3 | | | |
| | P | d | P | d | P | d | P | d |
| Δe45-53 to Δe46-54 | 0.57 | 0.50 | 0.00007 | 14 | 0.00008 | 14 | 0.00260 | 5.5 |
| Δe46-54 to Δe47-55 | 0.37 | 0.82 | 0.88 | 0.13 | 0.00001 | 26 | 0.00001 | 25 |
| Δe45-53 to Δe47-55 | 0.47 | 0.65 | 0.00007 | 14 | 0.00002 | 20 | 0.00069 | 14 |

### Supplemental Figures

#### Figure S1 - Equilibration During MD runs

A

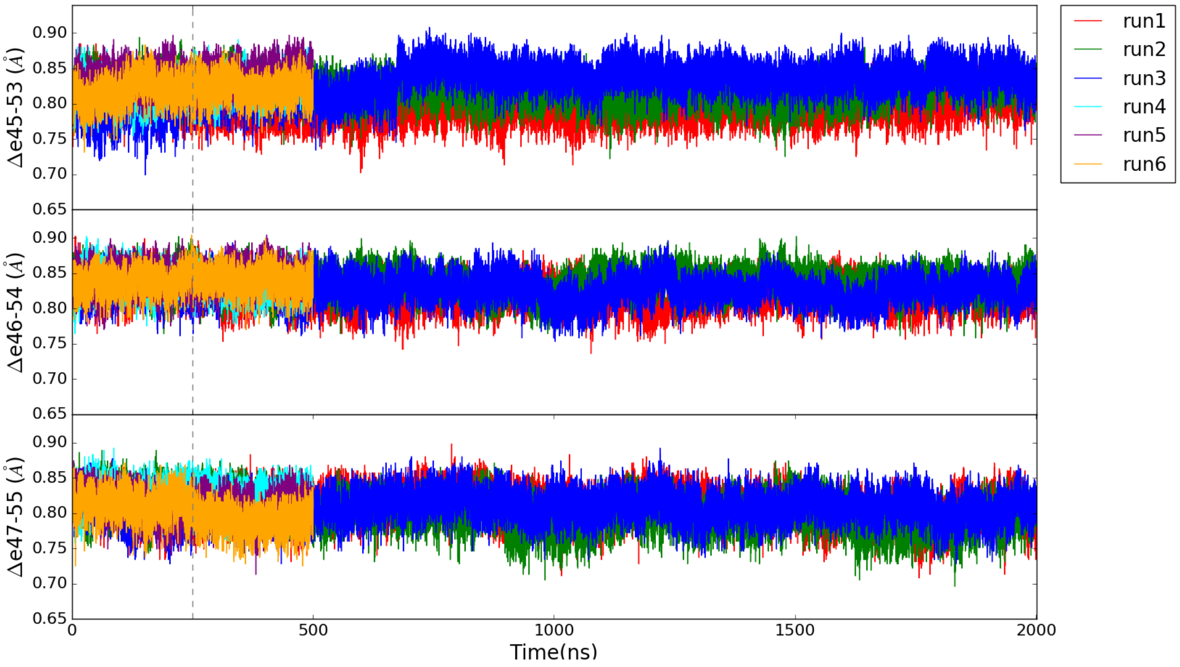


B

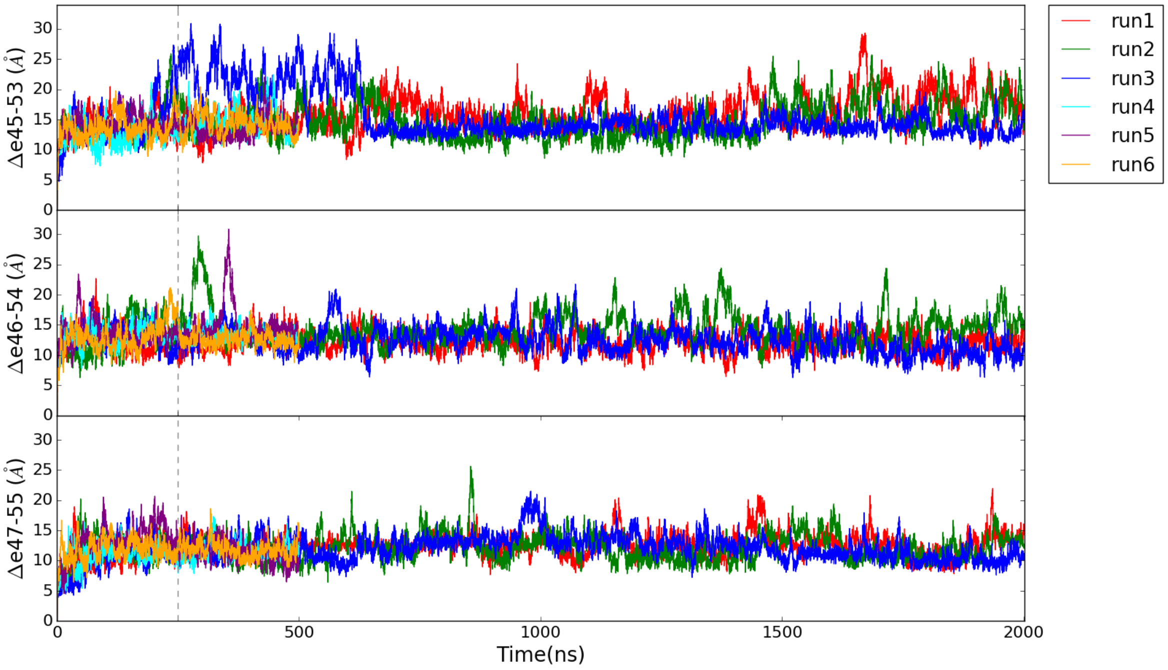


**Figure S1 Equilibration during MD runs.** In order to be sure our equilibration time of 250 nsec was sufficient, we assessed both helicity, **A**, as well as rmsd, **B** of the backbone from the starting conformation during both periods. The data before 250 usec, shown by the dotted line, was considered an equilibration period and not used for analysis. No systematic differences were seen after 250 usec, suggesting this was more than sufficient equilibration time.

#### Figure S2 – Empirical Alignment

```
                  A-helix             AB-loop              B-helix               BC-loop           C-helix
VMTEDMPLEISYVPSTYLTEITHVSQALLEVEQLLNAPDLCAKDFEDLFKQEESLKNIKDSLQQSSGRIDIIHSKKTAAL QSATPVERVKLQEALSQLDFQWEKVNKMYKDRQG                Δe4
VMTEDMPLEISYVPSTYLTEITHVSQALLEVEQLLNAPDLCAKDFEDLFKQEESLKNIKDSLQQSSGRIDIIHSKKTAAL QSATPVERVKLQEALSQLDFQWEKVNKMYKDRQG                Δe4
VMTEDMPLEISYVPSTYLTEITHVSQALLEVEQLLNAPDLCAKDFEDLFKQEESLKNIKDSLQQSSGRIDIIHSKKTAAL QSATPVERVKLQEALSQLDFQWEKVNKMYKDRQG                Δe4

     RFDRSVEKWRRFHYDIKIFNQWLTEAEQFLRK  TQIPENWEHAKYKWYLKqlakdlrqwqtnvdvandlalkll rdysaddtrkvhmiteninaswrsihkrvsereaaleethrl        Δe
     RFDRSVEKWRRFHYDIKIFNQWLTEAEQFLRK  TQIPENWEHAKYKWYLKELQDGIGQRQTVVRTLNATGEEII QQSSKTDASILQEKLGSLNLRWQEVCKQLSDRKKrvsereaaleethrl Δe4
     RFDRSVEKWRRFHYDIKIFNQWLTEAEQFLRK  TQIPENWEHAKYKWYLKELQDGIGQRQTVVRTLNATGEEII QQSSKTDASILQEKLGSLNLRWQEVCKQLSDRKKR               Δe4

      lqqfpldlekflawlteaettanvlqdatrke rlledskgvkelmkqwqdlqgeieahtdvyhnldensqkilrslegsddavllqrrldnmnfkwselrkkslnirshleassdqwkrl    Δe4
      lqqfpldlekflawlteaettanvlqdatrke rlledskgvkelmkqwqdlqgeieahtdvyhnldensqkilrslegsddavllqrrldnmnfkwselrkkslnirshleassdqwkrl    Δe4
      LEEQKNILSEFQRDLNEFVLWLEEADNIASIP LEPGKEQQLKEKLEQVKdlqgeieahtdvyhnldensqkilrslegsddavllqrrldnmnfkwselrkkslnirshleassdqwkrl    Δe4
      *      *  *   *  *   *  *   *                   *  *   *      *  *      *            *   *  *   *  *   *  *   *
      AbcDefgAbcDefgAbcDefgAbcDefgAbc          AbcDefgAbcDefgAbcDefgAbcDefgAbcDef       AbcDefgAbcDefgAbcDefgAbcDefgAbc
             A                A                       B                       B            C                 C
                  A-helix               AB-loop            B-helix               BC-loop           C-helix
```

Figure S2 Empirical identification of helices, Junctions and Heptad Repeats.

The primary sequences of the three targets in the D16:22 family context are shown, with the residues exhibiting >80% dynamic helicity highlighted in yellow. The location shaded in light blue near the N-terminal end of helix B of STR 3 is the triad consisting of {W,W,V} that is discussed in the MMGBSA energy analysis. The edit junction occurs at the transition between UPPERCASE and lowercase letters. These were aligned to the `AbcDefg` amphipathic bundled-helix heptad pattern, where hydrophobic residues occupy the `A` and `D` positions and are sequestered to the interior of the bundles. Brown asterisks highlight such positions that are both helical and hydrophobic in all three targets and all three STRs; lighter grey asterisks highlight positions that are helical and hydrophobic in at least one but not all STRs. Below that, in blue, an outermost A-D pair (brown residues in Figure 6) was selected for determining an STR vector in order to quantify STR bending angles as shown in Figure 9.

**Figure S3 – Experimental and Model Helicity**

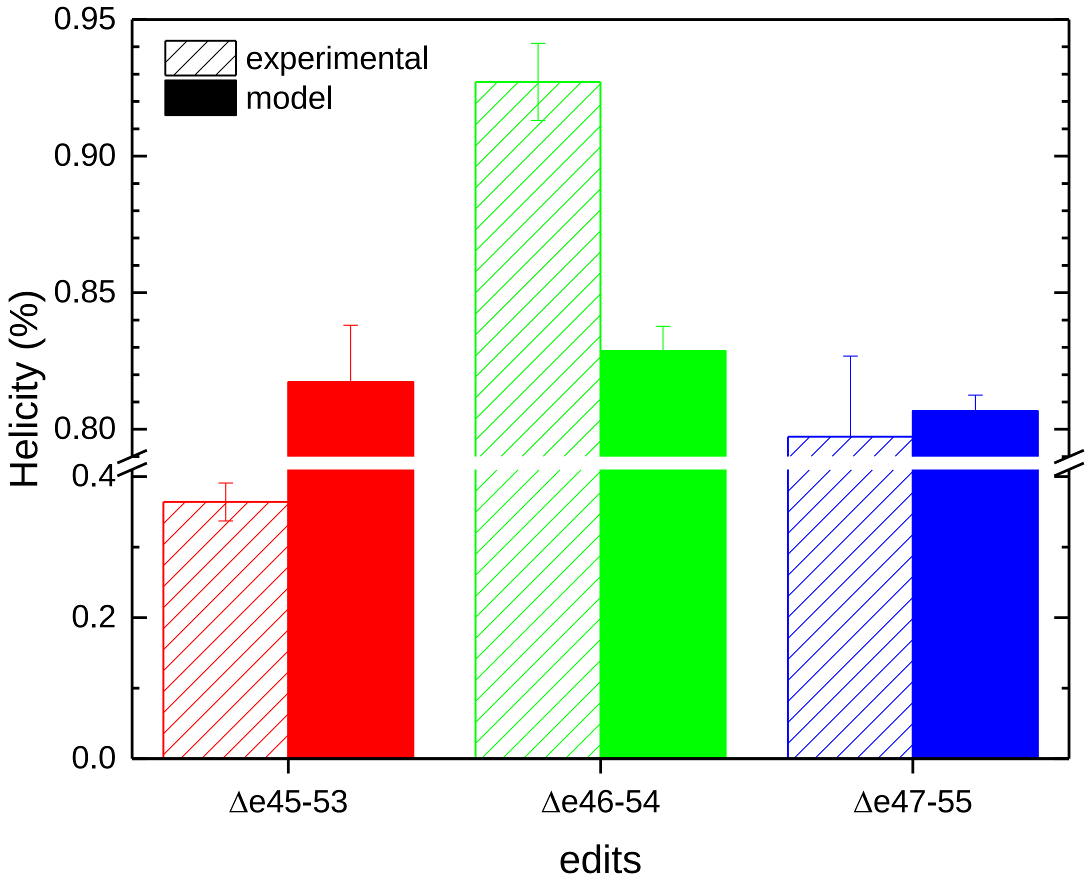


**Figure S3. Experiment vs Model Helicity Concordance.** Model helicity was assessed after modelled structures were relaxed and equilibrated to 300K as described. This was done independently for six simulation runs, and mean and standard error is shown. While the same rank order is reproduced, the differences between edits are not nearly as great as observed experimentally. This is most severe in the experimentally most perturbed structure, Δe45-53 becoming over predicted as helical in the modelled structure.

**Figure S4 – Location of Junction unfolding**

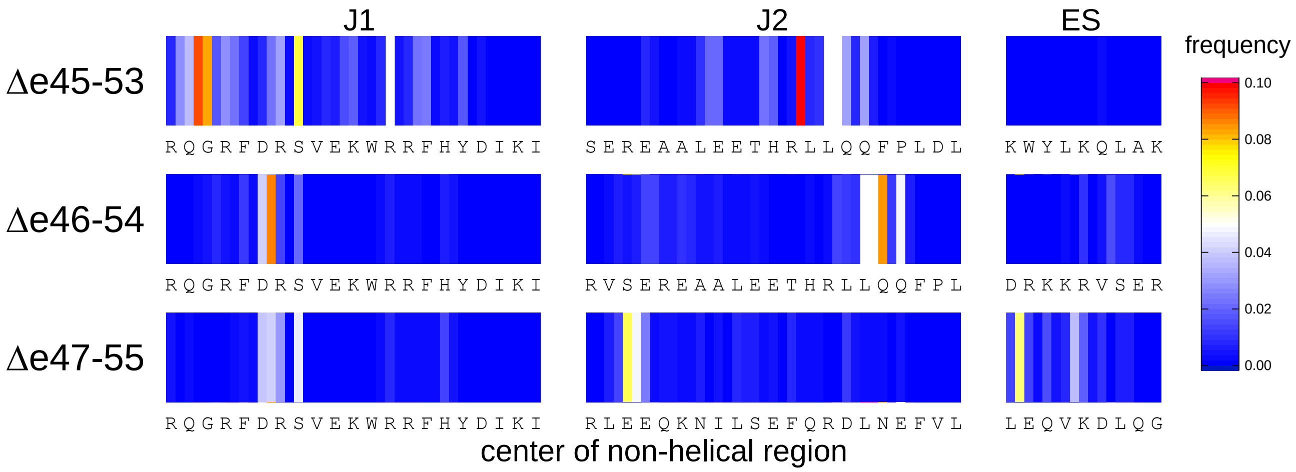


**Figure S4. Location of Unfolding**. Heatmaps of the location and frequency of the center of all detected non-helical regions in J1, J2 and ES are shown. We see that even though J1 has an identical primary sequence in all three targets, since it is before all edit sites, the location of unfolding is different, especially in the highly perturbed Δe44-53. The J2 region has a different sequence in all case since it occurs after the edit in two of the three cases, so it is perhaps not as surprising that the unfolding occurs in different relative locations. For J1 of Δe44-53 and J2 of Δe47-55, multiple distinct focal points for unfolding were observed. For the ES sites, only Δe47-55 had significant unfolding as shown in the main Figures 7 and 8.

**Figure S5 – Interpretation of STR Motional Correlation Maps**

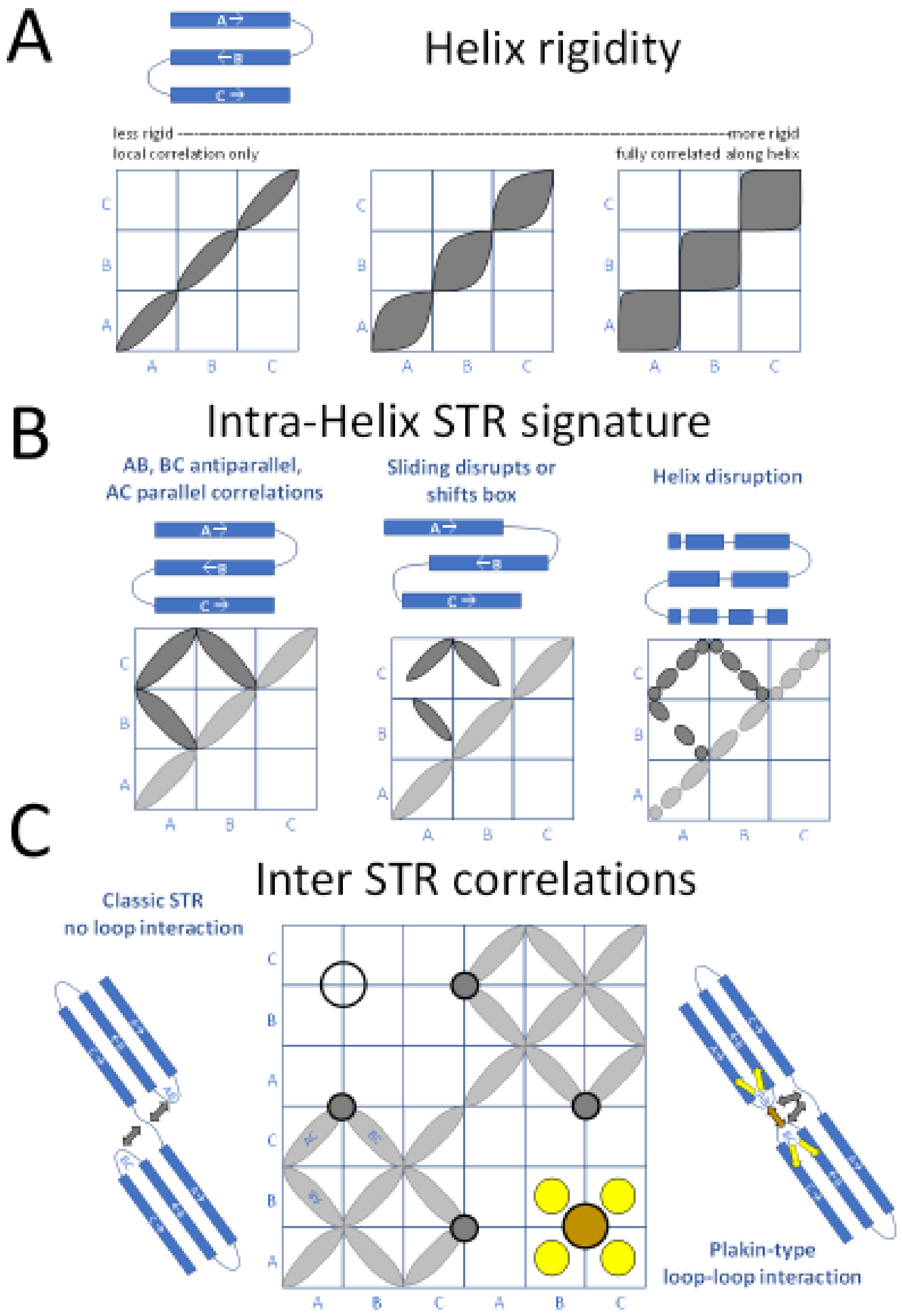

**Figure S5  STR correlation/contact signature interpretation**. **A** At the lowest level, motional correlation identifies secondary structural elements that act as rigid bodies. For helices, the more rigid they are the more fully the ends of the helix correlate with each other and the thicker, and more square, the helix signature gets; conversely thin helix traces are indicative of flexible elements. **B** Tertiary structure information can be seen by interaction between adjacent secondary structure elements. Here, the triple α-helical bundles manifests as a characteristic "STR box" structure. Weakening of bundling interactions resulting in independent (uncorrelated) motion will shift and weaken this signature. Similarly, disruption of the helices will break apart the box into smaller fragments. **C** Inter-STR interactions result in off-diagonal interactions. In particular a "plakin type" junction where there are direct interactions between the AB- and BC-loops of adjacent STR is easy to spot, and is shown in **gold**. Since the loops then link back the helices, four additional secondary interactions, **yellow**, can sometimes be seen as well. In contrast, the "classic STR" type junction, where the AB- and BC loops lie on opposite sides of the junction helix, results in no direct interaction and so no off-diagonal signals. Interactions between the junction helix and the loops, **grey**, occur in both cases, but are not well resolved from the general STR box signature.

**Figure S6 - Individual STR Correlation Maps**

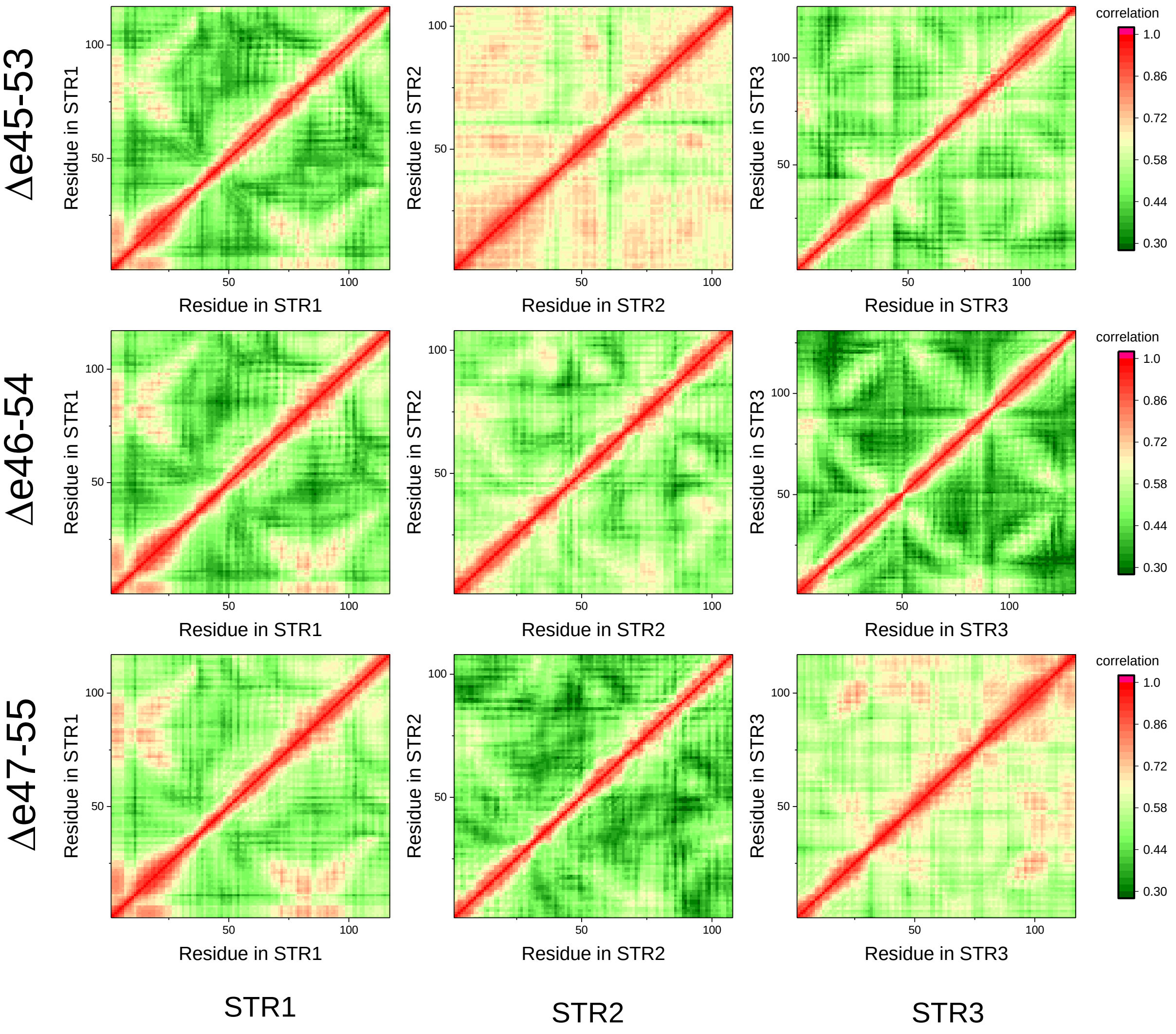


**Figure S6 Correlation analysis conducted independently on individual STRs**. This shows that excess rod flexibility of Δe45-53 was what was driving the abnormally high overall correlation of that edit. When conducted independently, overall correlation dropped and became more uniform and the "STR box pattern" was evident in most STRs – with the notable exception of the second STR of Δe45-53 and the third STR of Δe47-55. These are the two hybrid STRs, with internal edit sites and so composed of regions of two different wild-type STRs. (For Δe46-54, the edit site is close to the STR2-3 junction so all STRs are mostly wildtype). Both of these display elevated overall correlation indicative of excess motion, and also do not exhibit the characteristic "STR box" signature.

**Figure S7 – W replaced by V in De47-55 in heptad triad of STR 3**

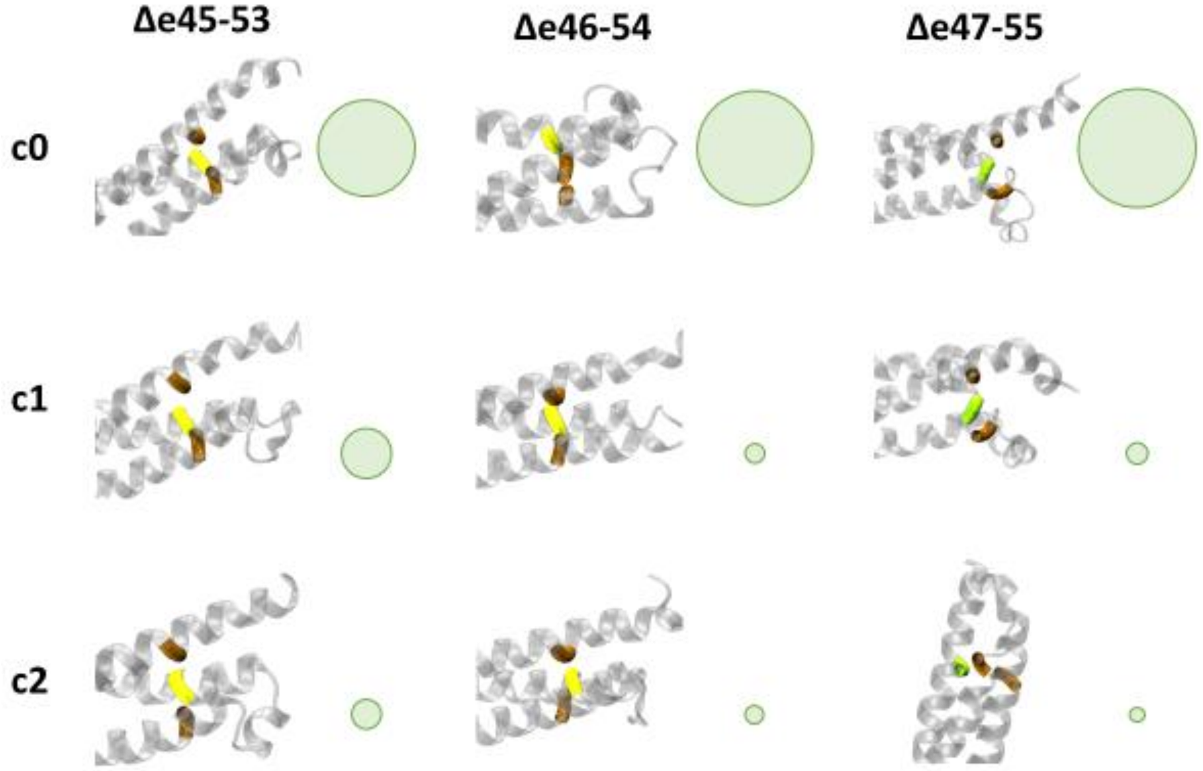


**Figure S7. The WWV triad of STR 3**. In the heptad triads at the end of STR3, a W, yellow, occurs in Δe45-53 and Δe46-54, but is replaced by a V, green, in Δe47-55. Their heptad hydrophobic partners are shown in brown. This region partially unbundles in Δe45-53, as shown by the 3 most abundant representative structures after clusters analysis, with green circles having areas proportional to cluster abundance. The dominant conformers of Δe47-55 have this end STR3 opened up.

**Figure S8 – Explicit Solvent RMSD evolution**

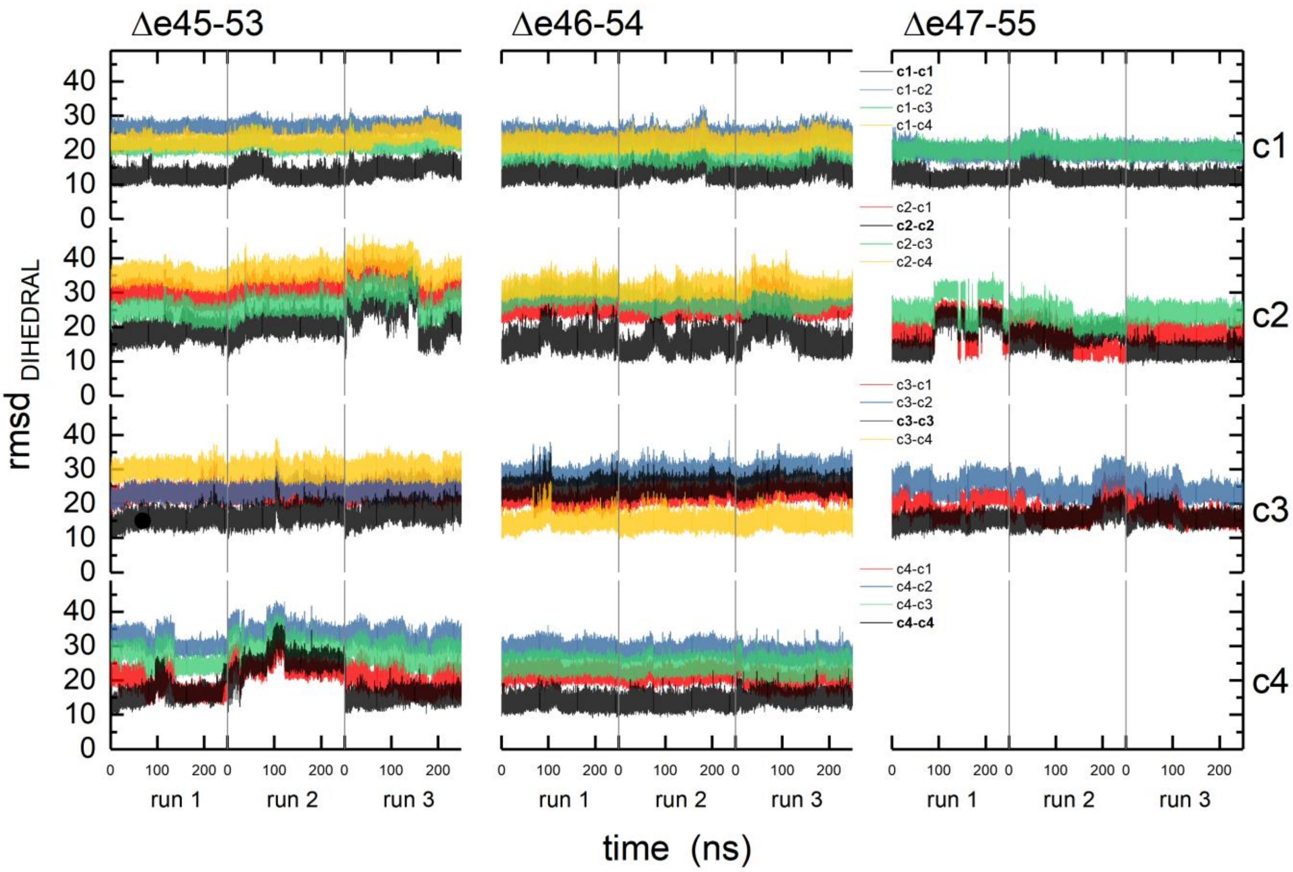


**Figure S8 - Evolution of explicit MD runs.** Explicit solvent MD runs were started from the representative conformers of the top 4 or 3 clusters (i.e. those with abundance > 1.5%). The RMSD distances of each run to all clusters as function of time are shown along each trajectory. The distance from its own starting point cluster is shown in **black**, and from heterologous clusters in colors as indicated. We can see that in all but one case the runs stayed closer to their own starting point structure, and do not rapidly evolve into another cluster. The sole exception was cluster 3 of Δe46-53 (a low abundance cluster, 2%) which appeared to rapidly move into a conformer similar to cluster 1 or 2 of that target during equilibration, and stay there during the run.

**Figure S9 – Explicit/Implicit helicity concordance by residue**

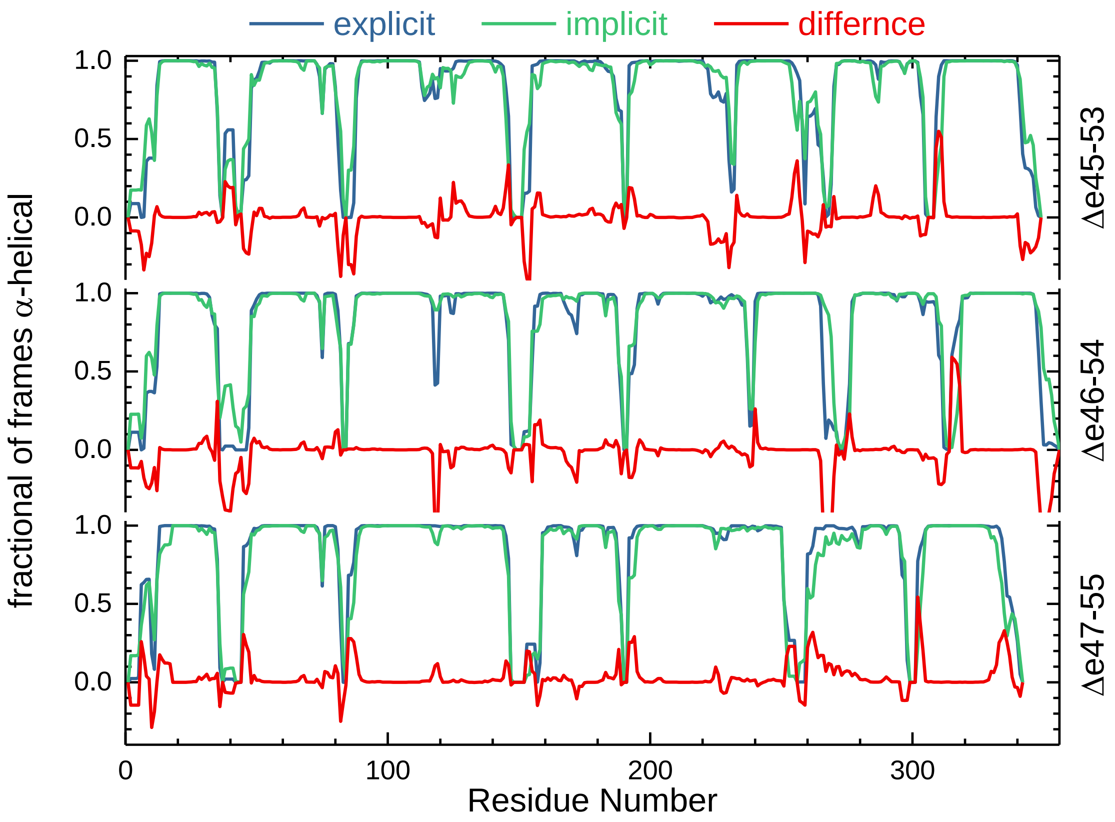


**Figure S10 comparison of helicity by residue for implicit and explicit runs**. Explicit and implicit runs show the same helicity pattern; where they differ chiefly occurs at the edges of the helicities which may differ by only a few residues on average.

**Figure S10 – Explicit Solvent helicity**

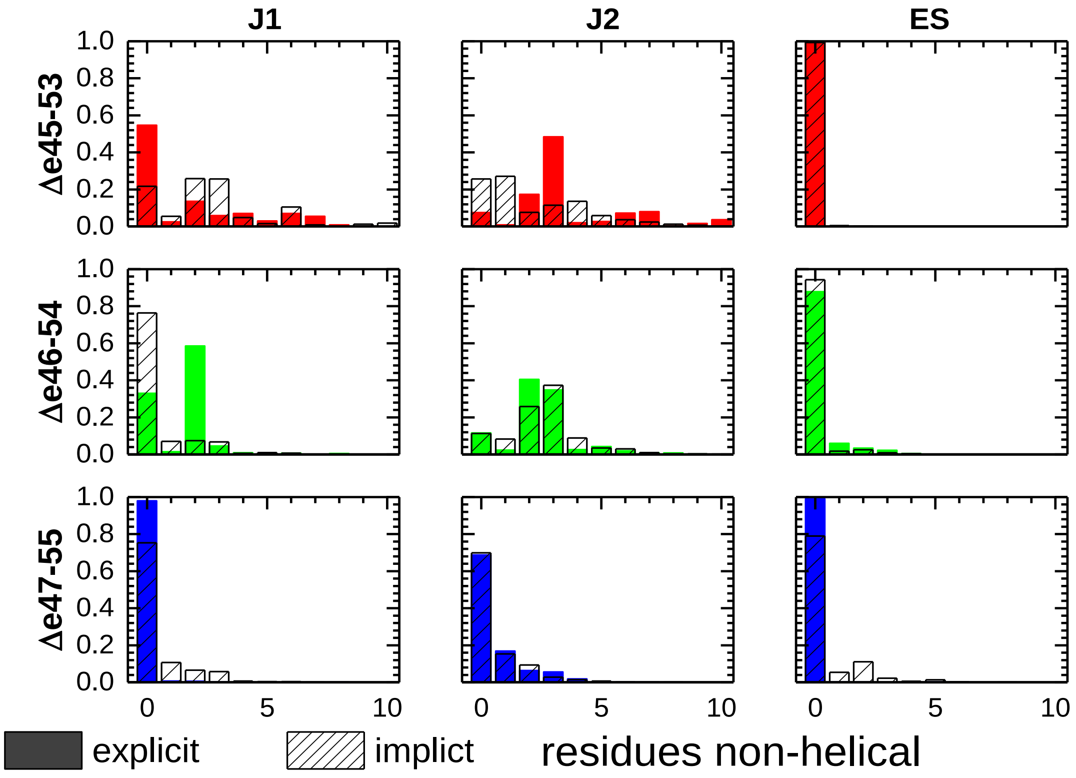


**Figure S10 Concordance of Explicit vs Implicit Runs with Respect to Helicity**. In general, the same basic features were observed, in that the ES sites were only minimally impacted, demonstrating that edits act distally, not locally; and both junction regions of Δe45-53 were significantly unfolded as in implicit solvent runs. The J2 regions of Δe46-54 and Δe47-55 are concordant with implicit, and J1 regions are once again more folded that J2. This supports the main contention of this work, that edits do not act to perturb structure by local effects at ES sites, but rather act distally, at the junctions.

**Figure S11 – Explicit Solvent Bending**

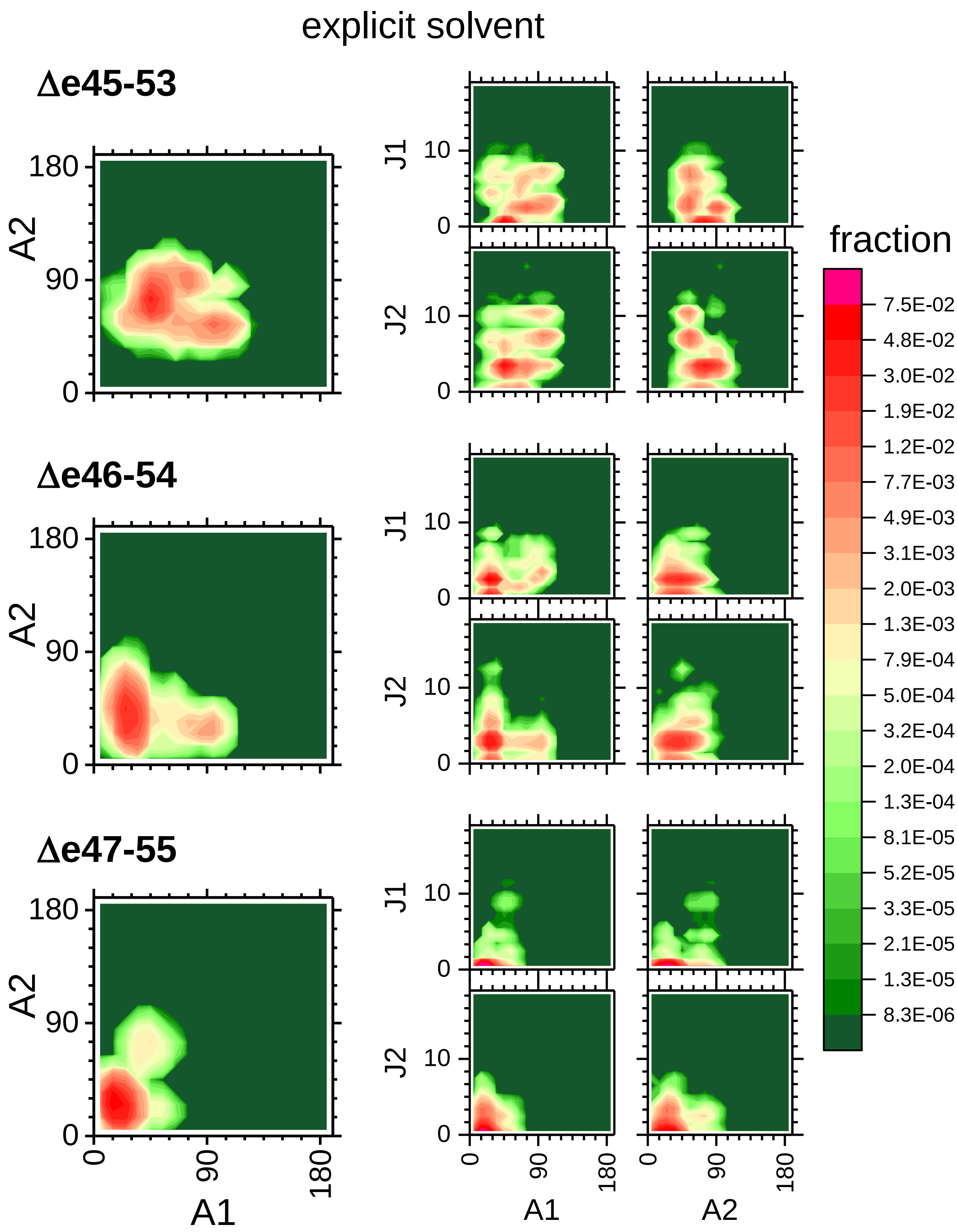

**Figure S11  Bending Analysis of the Explicit Solvent Data Set**. Explicit MD simulations were analyzed in the same fashion as the implicit data, and this figure is homologous to Figure 9. While the data is less robust due to lower sampling, the same maps look similar and the same general features are observed:

- for Δe45-53, a second population with elevated A1 is seen (A1 vs A2 graph) as in implicit.
- Increased bending at A1 is correlated with increased J1 unwinding (i.e. density in a diagonal up and to the right), especially for Δe45-53.
- The Δe47-55 edit also exhibits lower bending angles and junction unwinding, as in implicit runs.

### References

Åqvist, J., Wennerström, P., Nervall, M., Bjelic, S., and Brandsdal, B.O. (2004). Molecular dynamics simulations of water and biomolecules with a Monte Carlo constant pressure algorithm. Chem. Phys. Lett. *384*, 288–294.

Chowdhury, R., Rasheed, M., Keidel, D., Moussalem, M., Olson, A., Sanner, M., and Bajaj, C. (2013). Protein-protein docking with F(2)Dock 2.0 and GB-rerank. PloS One *8*, e51307.

Essmann, U., Perera, L., Berkowitz, M.L., Darden, T., Lee, H., and Pedersen, L.G. (1995). A smooth particle mesh Ewald method. J. Chem. Phys. *103*, 8577–8593.

Gibson, D.G., Glass, J.I., Lartigue, C., Noskov, V.N., Chuang, R.-Y., Algire, M.A., Benders, G.A., Montague, M.G., Ma, L., Moodie, M.M., et al. (2010). Creation of a bacterial cell controlled by a chemically synthesized genome. Science *329*, 52–56.

Jorgensen, W.L., Chandrasekhar, J., Madura, J.D., Impey, R.W., and Klein, M.L. (1983). Comparison of simple potential functions for simulating liquid water. J. Chem. Phys. *79*, 926–935.

Kabsch, W., and Sander, C. (1983). Dictionary of protein secondary structure: pattern recognition of hydrogen-bonded and geometrical features. Biopolymers *22*, 2577–2637.

Koenig, M., Monaco, A.P., and Kunkel, L.M. (1988). The complete sequence of dystrophin predicts a rod-shaped cytoskeletal protein. Cell *53*, 219–228.

Miller, B.R., McGee, T.D., Swails, J.M., Homeyer, N., Gohlke, H., and Roitberg, A.E. (2012). *MMPBSA.py* : An Efficient Program for End-State Free Energy Calculations. J. Chem. Theory Comput. *8*, 3314–3321.

Mongan, J., Simmerling, C., McCammon, J.A., Case, D.A., and Onufriev, A. (2007). Generalized Born Model with a Simple, Robust Molecular Volume Correction. J. Chem. Theory Comput. *3*, 156–169.

Sahni, N., Mangat, K., Le Rumeur, E., and Menhart, N. (2012). Exon edited dystrophin rods in the hinge 3 region. Biochim. Biophys. Acta *1824*, 1080–1089.

Winder, S.J., Gibson, T.J., and Kendrick-Jones, J. (1995). Dystrophin and utrophin: the missing links! FEBS Lett. *369*, 27–33.
